# Metatranscriptomic analysis reveals toxin-antitoxin system shifts in caries-associated oral microbiomes

**DOI:** 10.1101/2025.08.07.669164

**Authors:** Shri Vishalini Rajaram, Priyanka Singh, Erliang Zeng

## Abstract

**Background:** Toxin-antitoxin (TA) systems are bacterial regulatory elements involved in persistence, dormancy, and biofilm stability under stress conditions such as limited nutrients, acid exposure, and antimicrobial treatment. Although their roles are well-characterized in isolated pathogens, TA systems remain largely unexamined in the context of the oral microbiome. Given their potential involvement in microbial adaptation during oral diseases and treatment, we aimed to identify transcriptionally active TA systems and evaluate their condition-specific expression patterns across oral health states using metatranscriptomic data.

**Results:** We re-analyzed two publicly available supragingival metatranscriptomic datasets: a longitudinal dataset by Carda-Diéguez et.al. (Dieguez) capturing treatment transitions (fluoride (Fl), fluoride-arginine (Fl-Ar)) in caries-active and caries-free individuals, and a cross-sectional dataset by Ev et.al., (Ev) profiling caries-active lesions (CA), caries-inactive (CI), and caries-free (CF) states. After quality control, host sequences removal, rRNA removal, and HUMAnN3-based functional profiling, we identified 1,189 unique UniRef90 gene clusters with known TA associations. Differential analysis using ANCOM-BC revealed 197 TA-related gene clusters as significantly modulated in at least one condition. Of these, 22 gene clusters were differentially expressed in Dieguez, 38 in Ev, with 77 shared across both datasets. Distinct TA expression signatures were observed across healthy (CF, CI, baseline), lesion types (CAa, CAi, CAs), and treatment stages (Fl, Fl-Ar), suggesting condition-specific regulatory activity. Functional annotation of differentially expressed TA genes using the Toxin-antitoxin system database, eggNOG-mapper, and InterProScan, revealed 18 genes with strong evidence for their involvement in key bacterial stress-response pathways. These included members of the ParD-ParE, RelE/StbE-RelB/StbD, FitA, and YafQ families, which are annotated with mRNA catabolism, transcriptional repression, and prokaryotic defense-associated pathways.

**Conclusion:** This is the first metatranscriptomic analysis profiling functional shifts in toxin-antitoxin systems across oral healthy, caries, and treatment states. TA systems showed dynamic and condition-specific expression, with pathway annotations suggesting their roles in microbial persistence, stress adaptation, and ecological remodeling during caries progression. These findings open new avenues for targeting microbial stress modules in precision microbiome therapeutics.

## Background

Recent epidemiological studies indicate that untreated dental caries affects approximately 3.5 billion adults and 572 million children globally [1, 2]. Caries is a multifactorial disease caused by the interplay of diet, genetics, microbes, and host factors over time. While sugar intake and hygiene behaviors are established contributors, the past two decades have spotlighted the oral microbiome as a central driver of caries. [3, 4]. The oral microbiome is composed of a complex polymicrobial biofilm that plays a central role in caries development. High sugar intake and repeated low pH episodes select aciduric bacteria, resulting in dysbiosis dominated by species such as *Streptococcus mutans*, *Lactobacillus*, *Actinomyces*, and *Scardovia*. This microbial imbalance, characterized by dysbiotic flora, drives net mineral loss from teeth. Typically, a healthy plaque community is maintained in balance by a complex web of interspecies interactions and the buffering capacity of saliva. However, the homeostatic balance can be disrupted by environmental stresses such as frequent acidification, oxidative stress, or nutrient deprivation [5]. Microbial interactions, both cooperative and competitive, further modulate the composition and pathogenicity of the community. The persistence of acidogenic, acid-tolerant bacterial communities at the tooth surface ultimately leads to carious lesions.

From a general microbiological perspective, a bacterium can employ numerous adaptive mechanisms to survive stress and harsh conditions. General stress-response pathways, and two-component regulatory systems are activated to deal with the stress [6, 7]. One such system, still understudied in the context of the oral microbiome, is the toxin-antitoxin (TA) system. Bacterial TA systems are genetic modules consisting of a stable toxin and a labile antitoxin encoded together in an operon [8–10].

The toxin (usually a protein) can inhibit essential cellular processes, causing growth arrest or cell death, while the antitoxin (protein or small non-coding RNA) neutralizes the toxin under normal conditions. Currently, TA systems are classified into eight types (I-VIII), based on the nature and mechanism of the antitoxin. These eight systems serve diverse functions in cellular physiology and survival, acting as switches that help bacteria adapt to adverse conditions [10, 11]. Under stressors such as nutrient starvation, antibiotics, or DNA damage, antitoxins degrade, releasing free toxins that induce reversible dormancy. This mechanism leads to the formation of persister cells, phenotypic variants that exhibit high tolerance to antibiotics and contribute to long-term bacterial survival [8, 11]. TA systems can also defend against bacteriophage attack by triggering abortive infection, sacrificing infected cells to protect the overall community [11, 12]. Moreover, many pathogenic bacteria carry numerous TA loci that aid in their adaptation to host-induced stresses (oxidative stress, immune responses) and thus enhance colonization and virulence [9, 13, 14]. While TA systems are now recognized as integral to bacterial stress physiology, their roles in complex microbial communities remain underexplored [9, 15]. In the microbiome context, TA systems may modulate both intra- and inter-species dynamics, including interactions with viruses, fungi, and archaea.

TA modules have been characterized in oral species like *Streptococcus mutans*, including type I, II, and tripartite systems involved in stress tolerance and persistence [16–19], supporting their relevance to oral microbial ecology. Thus, TA systems might be an unrecognized factor in the recalcitrance of cariogenic biofilms and the difficulty of entirely eradicating the disease.

Despite the known roles of TA systems in bacterial stress survival and pathogenesis, very few studies have examined these systems in the context of oral health and disease. Most research on TA systems has focused on model organisms such as *Escherichia coli* and pathogens causing systemic infections [9, 14, 15]. Although recent investigations have begun to characterize TA modules in oral bacteria [16–19], there remains a lack of evidence directly linking TA system activity to the onset or progression of oral diseases such as dental caries. This represents a critical knowledge gap: we do not yet know whether TA-mediated stress responses in oral microbes tangibly contribute to the cariogenic shift and tooth decay in vivo. Given the ubiquity of TA systems and their impact on key bacterial behaviors (stress tolerance, biofilm formation, persistence), it is essential to determine if and how they influence the balance of the oral microbiome. Typically, in a healthy biofilm, a balance between acid-producing and alkali-producing bacteria, or those less acid-tolerant, prevents extreme pH drops. If certain bacteria deploy TA-mediated tactics to eliminate competitors or secure nutrients, this could disrupt the equilibrium.

In this direction, we hypothesize that bacteria in caries-associated microbiomes leverage TA systems to survive chronic environmental stress, promote persistence, and shape community dynamics. As we have evidenced earlier, the chronic dental caries, characterized by an acidic environment, oxidative stress, and the survival of several bacterial populations, could be a valid reason to connect this to the TA systems. Another way TA systems could contribute to caries is through modulating interspecies competition and biofilm stability. Therefore, we posit that bacteria in the microbiome utilize TA systems to ensure survival, facilitate stress responses, and promote persistence against stressors. **Figure *1***. Our research focuses on a comprehensive analysis of the metatranscriptomic datasets to assess the presence and functional contribution of TA systems in the microbiome of caries-free, caries-affected and caries-treated patients. Such insights could fundamentally improve our understanding of caries etiology, potentially revealing novel molecular targets for preventive or therapeutic strategies.

**Figure 1.**
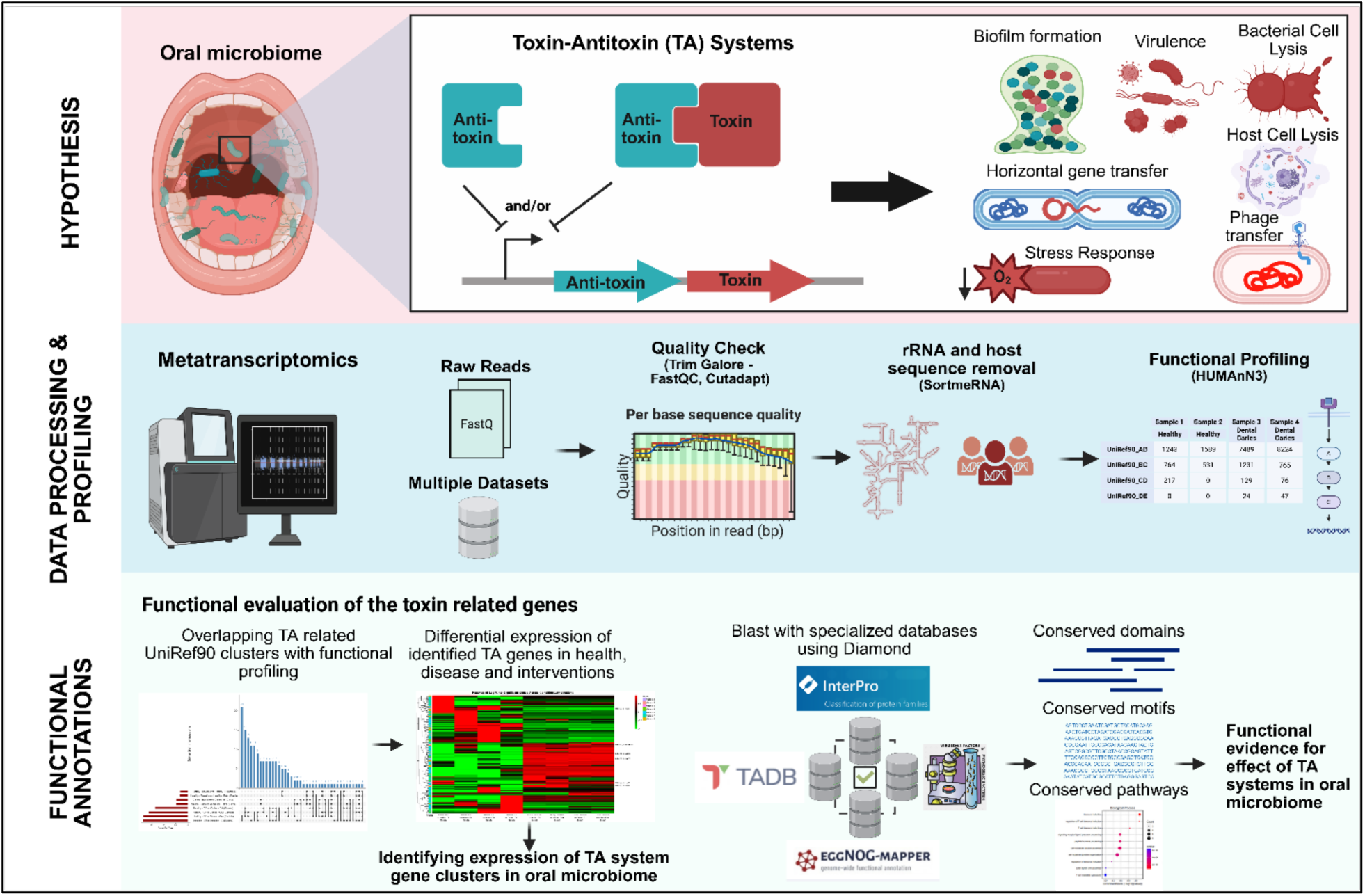
Workflow for identifying and functionally validating toxin-antitoxin (TA) systems in oral microbiome. Publicly available metatranscriptomic datasets from healthy, caries-affected, and treated plaque samples were reanalyzed. After quality control, host and rRNA removal, HUMAnN3 was used to quantify UniRef90 gene families. TA-associated clusters were identified by sequence homology against specialized databases. Conserved domains, motifs, and pathways were then annotated to infer functional roles related to microbial stress response and caries-associated dysbiosis.

## Methods

### Data Acquisition

Metatranscriptomic datasets were obtained from the National Center for Biotechnological Information - Sequence Read Archive (NCBI-SRA), specifically the datasets from Carda Dieguez et al. (2022) [20] and Ev et. al. (2023) [21]. Hereafter, we refer to these datasets as the Dieguez and Ev datasets. Both datasets include supragingival biofilm samples collected from individuals representing different oral health states. The Dieguez dataset consists of raw FASTQ reads generated on the Illumina NextSeq 550 platform (2 × 150 bp reads), whereas the Ev dataset was sequenced on the Illumina HiSeq 3000 (2 × 150 bp reads).

Despite both datasets examining the oral microbiota, they differ in study design and participant classification. Sample collection protocols varied with the Dieguez dataset focusing on overall supragingival plaque, while the Ev dataset targeted distinct lesion sites within each oral health classification. The Dieguez dataset comprises longitudinal sequencing data from 18 caries-active and 20 caries-free patients, with samples collected at baseline (week 1), after fluoride treatment (3 months), and following fluoride plus arginine dentifrice treatment (6 months). The Ev dataset employs a cross-sectional design, categorizing participants into groups of caries-active (CA, n = 5), caries-inactive (CI, n = 3), and caries-free (CF, n = 6) based on the decayed, missing, and filled teeth/surfaces (DMF-T/S) index. Three localizations (Active non-cavitated caries lesions (ANCL) - CAa, Inactive non-cavitated caries lesions (INCL) - CAi, and sound surfaces CAs) were sampled from each caries-active patient. The complementary study designs of these datasets offer the opportunity to explore both temporal (Dieguez) and spatial (Ev) variations in caries-associated microbiomes **(*Table 1*)**.

**Table 1.** Description of the Dieguez (longitudinal) and Ev (cross-sectional) datasets used for TA system analysis. The samples in the Dieguez dataset were collected from Healthy and Caries individuals who were given 1450 ppm of Fl as Sodium monofluorophosphate with insoluble calcium compound – for one week 1 week (Baseline), for 3 months (Fl) and for 9 months (Fl-Ar) (Fl + 1.5% arginine).

|  | Dieguez - Dataset [20] | Ev - Dataset [21] |
| --- | --- | --- |
| **Data type** | Longitudinal | Cross-sectional |
| **Groups/Condition (Sample size)** | Caries-active (Caries, 26)<br>Baseline (7)<br>Fluoride (FI, 11)<br>Fluoride-Arginine (FI-Ar, 17)<br>Caries-free (Healthy, 27)<br>Baseline (6)<br>Fluoride (FI, 17)<br>Fluoride-Arginine (FI-Ar, 18) | Caries-active (CA) (Caries, 5)<br>CAa (ANCL sites)<br>CAi (INCL sites)<br>CAs (sound dental surfaces)<br>Caries-inactive (CI) (Healthy, 3)<br>Caries-free (CF) (Healthy, 6) |
| **Sample localization** | Supragingival plaque | Supragingival plaque |
| **Sequencing depth** | 2 * 150 bp (paired end) | 2 * 150 bp (paired end) |
| **Sequencing type** | Illumina Miseq/NextSeq 550 | Illumina HiSeq 3000 |

\*ANCL - Active non-cavitated caries lesions, INCL - Inactive non-cavitated caries lesions

### Data Preprocessing

Raw sequence data were processed using a custom computational pipeline optimized for high throughput metatranscriptomic analysis. Quality control procedures included the assessment of sequencing quality (FastQC (v0.12.1)) and trimming of low-quality bases and adapter sequences (Cutadapt (v2.10)) using Trim Galore (v0.6.10) [22]. To eliminate non-target sequences, ribosomal RNA (rRNA) and host-derived reads were filtered out using SortMeRNA (v4.3.7) [23] against reference databases, including SILVA [24], Greengenes [25], and the GRCh38 human genome. Filtered reads were retained for downstream functional profiling.

### Functional Profiling

We profiled the datasets using HUMAnN3 (v3.9) [26], which mapped gene families and metabolic pathways to the UniRef90 protein cluster database. The functional profiling provided specific gene families and metabolic pathways across different health, affected, and treatment conditions. Gene and pathway expression tables were obtained from functional profiling. Gene expression tables contain identified UniRef90 clusters and their expression across samples. Toxin-antitoxin (TA) system-associated genes were identified by intersecting UniRef90 clusters from the expression tables with a curated list of TA-associated UniRef90 entries obtained from the UniRef database (https://www.uniprot.org/uniref, November 19, 2024) by querying “toxin” or “toxic*”.

### Differential Expression Analysis

From the HUMAnN3 functional gene tables, we identified UniRef90 clusters detected in the datasets and intersected them with a curated list of TA-associated UniRef90 entries. We analyzed their datasets separately because the Dieguez and Ev studies differ in experimental design and sampling. For each dataset, we retained only TA-associated genes; thus, the Dieguez analysis includes TA genes unique to Dieguez plus those shared with Ev, and vice versa. This approach enables direct comparison of TA expression profiles shared across studies versus those unique to each dataset.

Differential expression analysis was performed on these two modified gene tables using Analysis of Covariance in Microbiome data with Bias Correction (ANCOM-BC) [27]. We tested every pairwise contrast among healthy, caries-affected, and treated groups. In the Ev study, the healthy categories were Healthy-CF and Healthy-CI, and the caries categories were CAa, CAi, and CAs. In the Dieguez study, healthy and caries samples corresponded to their respective baseline sets, with additional treated groups Fl and Fl-Ar. The significant genes identified in these comparisons capture the biological distinctions between conditions and sharpen our interpretation of temporal and spatial variation.

### Functional Annotation of TA Systems

While differential expression analysis identifies statistically significant TA genes, functional annotation is critical to contextualize their biological relevance. For this purpose, the FASTA sequences of functionally profiled UniRef90 gene clusters were extracted from the UniRef90 database and aligned against the sequences from the Toxin-Antitoxin Database (TADB, v3.0) [28] and the Virulence Factor Database (VFDB) [29]. All alignments were performed using DIAMOND (v2.0.10) [30] with default parameters unless otherwise specified. TADB is the most comprehensive repository of the experimentally validated and computationally predicted bacterial TA system genes [28]. Aligning to these genes helped to refine the list of statistically significant TA system genes from differentially expressed gene list. We cross-referenced our significant TA genes with VFDB to find those that also encode virulence factors [29]. UniRef90 sequences were annotated with eggNOG-mapper v2.1.12 [31], using its built-in database and thresholds of ≥50 % identity and ≥80 % query coverage. EggNOG-mapper assigned Pfam families, COGs, and GO/KEGG pathway terms. Additional annotations came from InterProScan v5.72-103.0 [32], which integrates signatures of protein families (Pfam, PROSITE, FunFam), specific pathways or processes (GO, Reactome, MetaCyc), and domains (CDD - Conserved Domains Database). Together, these pipelines deliver high-sensitivity, high-specificity functional mapping of toxin-antitoxin genes and support mechanistic insight into their roles.

## Results

### Quality Control and Functional Profiling of Supragingival Microbiomes

The metatranscriptomic datasets from the Dieguez (longitudinal) and Ev (cross-sectional) studies of supragingival plaque were preprocessed. The initial raw read quality was assessed using FastQC, with most samples demonstrating high per-base sequence quality, minimal adapter contamination, and no major GC bias. Adapter trimming and quality filtering were performed using Trim Galore (Cutadapt), followed by the removal of host (human) and ribosomal RNA contaminants using SortMeRNA. Given the highest proportion of rRNA and host reads, only 3-15% of the total reads were retained for further analysis. The number of reads retained at each stage is summarized in ***Table 2*** with detailed information in the supplementary table **S1**. The retention of high-quality, non-contaminant reads enabled reliable downstream functional characterization. Functional annotation was performed using HUMAnN3, which integrates MetaPhlAn for taxonomic profiling and Bowtie2/DIAMOND for translated search against the UniRef90 database. Across all samples, HUMAnN3 identified 209,523 gene clusters in Dieguez and 150,553 in Ev, providing the foundation for further downstream analysis to identify TA system genes.

**Table 2.**
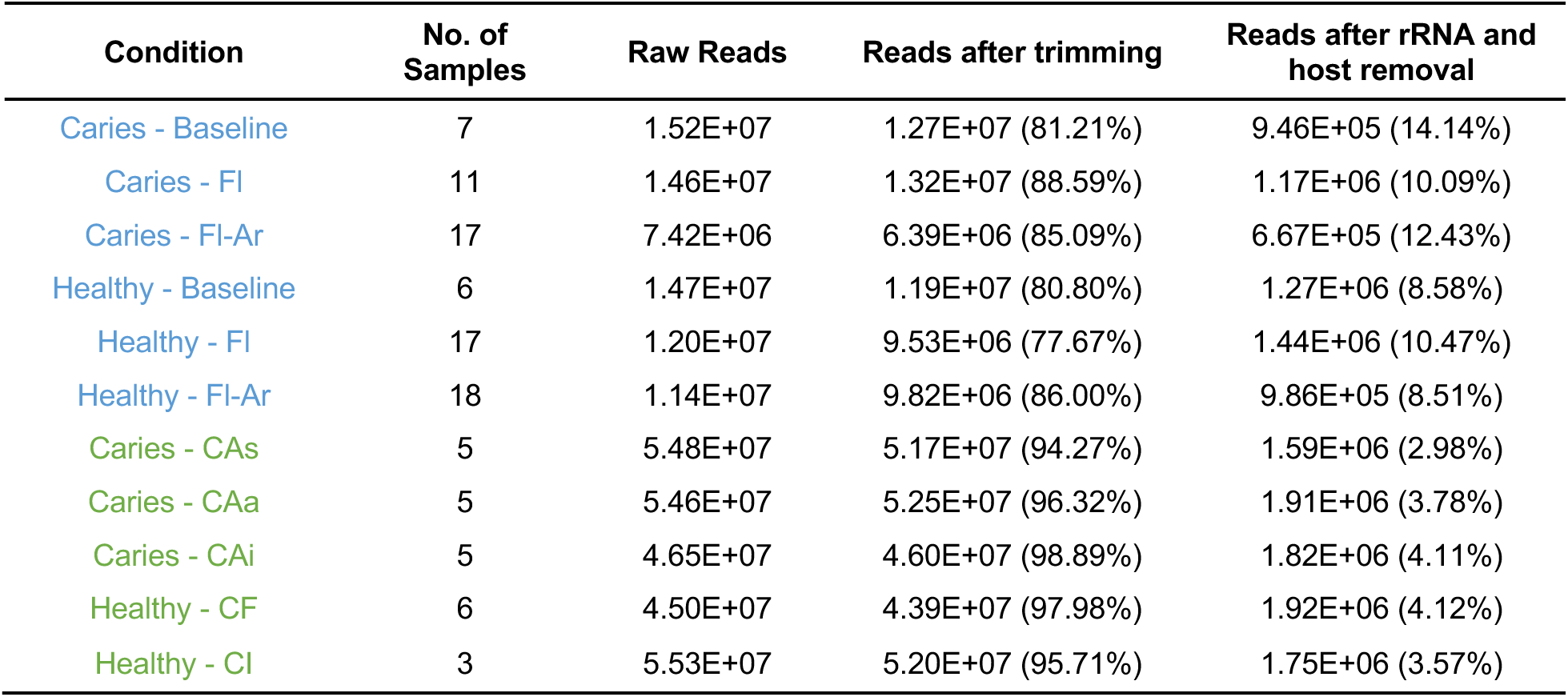
Summary of preprocessing and decontamination metrics for metatranscriptomic reads across all conditions. Read counts are reported as group-wise means. The “Reads after trimming” and “Reads after rRNA and host removal” columns both includes the number and the percentage of reads retained after adapter trimming and after removal of rRNA and host sequences. Conditions highlighted in blue and green correspond to datasets from Dieguez and Ev, respectively.

| Condition | No. of Samples | Raw Reads | Reads after trimming | Reads after rRNA and host removal |
| --- | --- | --- | --- | --- |
| Caries - Baseline | 7 | 1.52E+07 | 1.27E+07 (81.21%) | 9.46E+05 (14.14%) |
| Caries - FI | 11 | 1.46E+07 | 1.32E+07 (88.59%) | 1.17E+06 (10.09%) |
| Caries - FI-Ar | 17 | 7.42E+06 | 6.39E+06 (85.09%) | 6.67E+05 (12.43%) |
| Healthy - Baseline | 6 | 1.47E+07 | 1.19E+07 (80.80%) | 1.27E+06 (8.58%) |
| Healthy - FI | 17 | 1.20E+07 | 9.53E+06 (77.67%) | 1.44E+06 (10.47%) |
| Healthy - FI-Ar | 18 | 1.14E+07 | 9.82E+06 (86.00%) | 9.86E+05 (8.51%) |
| Caries - CAs | 5 | 5.48E+07 | 5.17E+07 (94.27%) | 1.59E+06 (2.98%) |
| Caries - CAa | 5 | 5.46E+07 | 5.25E+07 (96.32%) | 1.91E+06 (3.78%) |
| Caries - CAi | 5 | 4.65E+07 | 4.60E+07 (98.89%) | 1.82E+06 (4.11%) |
| Healthy - CF | 6 | 4.50E+07 | 4.39E+07 (97.98%) | 1.92E+06 (4.12%) |
| Healthy - CI | 3 | 5.53E+07 | 5.20E+07 (95.71%) | 1.75E+06 (3.57%) |

### Detection and Comparison of TA System Gene Clusters across Datasets

To investigate the role of TA systems in oral microbiome adaptation and stress regulation, we leveraged a curated reference of 919,926 TA-related UniRef90 gene clusters from the UniRef database. These were cross-mapped to the HUMAnN3-derived gene expression profiles. A total of 1015 TA gene clusters were detected in the Dieguez dataset, and 626 in the Ev dataset. Among these, 452 TA clusters were shared across both datasets (***Figure 2A***), forming the core supragingival TA system gene pool for downstream analysis.

**Figure 2.**
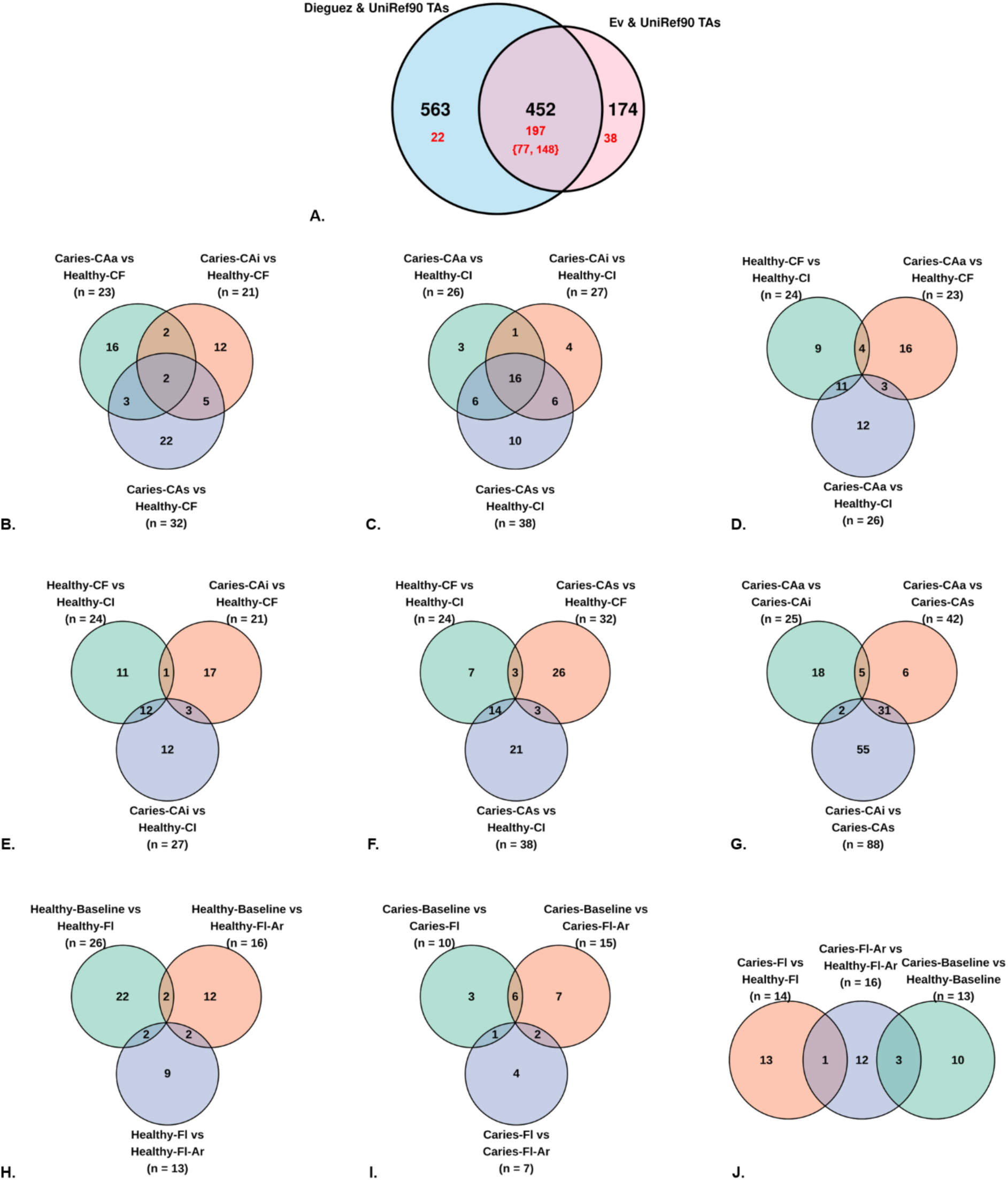
Comparison of TA system gene clusters across datasets and conditions. (A) Venn diagram shows overlap between TA-related UniRef90 clusters detected in the Dieguez and Ev datasets. Red numbers indicate differentially expressed genes identified using ANCOM-BC. (B-G) Condition-specific overlap of differentially expressed TA gene clusters in the Ev dataset across caries-active (CAa, CAi, CAs), caries-inactive (CI), and caries-free (CF) groups. Each panel shows shared and unique TA-related genes between two or three pairwise comparisons. (H-J) Condition-specific overlap of differentially expressed TA gene clusters in the Dieguez dataset across treatment stages (Baseline, Fl, Fl-Ar) within caries and healthy groups.

To begin with the downstream analysis, differentially expressed genes were evaluated using ANCOM-BC for all pairwise comparisons of health, disease, temporal, and spatial conditions in the Dieguez and Ev datasets **(Table 1**). Notably, 197 of the 452 shared clusters, when analyzed using ANCOM-BC, demonstrated statistically significant differential expression. Of these, 77 were significantly expressed in Dieguez, and 148 in Ev. Gene clusters that were unique to Dieguez and Ev and overlapped with the UniRef90 TA clusters also showed 22 and 38 significant genes, respectively **(*Figure 2A*)**. These overlaps suggest functional convergence across different experimental designs (longitudinal vs. cross-sectional) and emphasize that a subset of TA systems is consistently dysregulated in caries-associated conditions across spatial, temporal, and treatment gradients. The expression datasets for the overlapping entries are available in GitHub (https://github.com/biocoms/ta_systems_oral_mt/tree/main/uniref_tox_abundance), and the complete list of differentially expressed gene clusters is in the supplementary table **S2**.

### Baseline Variation in TA System Gene Expression within Healthy Individuals

At baseline, transcriptional comparisons between Healthy-CF and Healthy-CI supragingival samples identified 24 significantly differentially expressed TA gene clusters. Of these, 17 were upregulated in Healthy-CF and 7 in Healthy-CI, indicating divergent TA system activity between the two states (**Table 3**).

**Table 3.** Summary of down and up regulated significantly differentially expressed gene clusters from ANCOM-BC across contrasts.

| Group | Condition Comparisons | No. of Down Regulated Gene Clusters (%) | No. of Up Regulated Gene Clusters (%) |
| --- | --- | --- | --- |
| Healthy | Healthy-CF vs. Healthy-CI | 7 (29.17%) | 17 (70.83%) |
| Healthy_vs_Caries | Caries-CAa vs. Healthy-CF | 19 (82.61%) | 4 (17.39%) |
| Healthy_vs_Caries | Caries-CAa vs. Healthy-CI | 3 (11.54%) | 23 (88.46%) |
| Healthy_vs_Caries | Caries-CAi vs. Healthy-CF | 20 (95.24%) | 1 (4.76%) |
| Healthy_vs_Caries | Caries-CAi vs. Healthy-CI | 3 (11.11%) | 24 (88.89%) |
| Healthy_vs_Caries | Caries-CAs vs. Healthy-CF | 31 (96.88%) | 1 (3.13%) |
| Healthy_vs_Caries | Caries-CAs vs. Healthy-CI | 4 (10.53%) | 34 (89.47%) |
| Healthy_vs_Caries;<br>Timepoint_Healthy_vs_Caries | Caries-Baseline vs. Healthy-Baseline | 9 (69.23%) | 4 (30.77%) |
| Spatial_Caries | Caries-CAa vs. Caries-CAi | 3 (12.00%) | 22 (88.00%) |
| Spatial_Caries | Caries-CAa vs. Caries-CAs | 1 (2.38%) | 41 (97.62%) |
| Spatial_Caries | Caries-CAi vs. Caries-CAs | 1 (1.14%) | 87 (98.86%) |
| Timepoint_Caries | Caries-Baseline vs. Caries-FI | 4 (40.00%) | 6 (60.00%) |
| Timepoint_Caries | Caries-Baseline vs. Caries-FI-Ar | 3 (20.00%) | 12 (80.00%) |
| Timepoint_Caries | Caries-FI vs. Caries-FI-Ar | 2 (28.57%) | 5 (71.43%) |
| Timepoint_Healthy | Healthy-Baseline vs. Healthy-FI | 2 (7.69%) | 24 (92.31%) |
| Timepoint_Healthy | Healthy-Baseline vs. Healthy-FI-Ar | 14 (87.50%) | 2 (12.50%) |
| Timepoint_Healthy | Healthy-FI vs. Healthy-FI-Ar | 10 (76.92%) | 3 (23.08%) |
| Timepoint_Healthy_vs_Caries | Caries-FI vs. Healthy-FI | 12 (85.71%) | 2 (14.29%) |
| Timepoint_Healthy_vs_Caries | Caries-FI-Ar vs. Healthy-FI-Ar | 5 (31.25%) | 11 (68.75%) |

In Healthy-CF, transcripts corresponding to both toxin and antitoxin components were elevated compared to Healthy-CI. Upregulated antitoxins included eight RHH domain-containing gene clusters (UniRef90_C9MUM6, UniRef90_E7RXG0, UniRef90_L1NQ05, UniRef90_U1QP00, UniRef90_U2BYZ5, UniRef90_U2KZ56, UniRef90_U2LXT4, UniRef90_X8H3M8). Concurrently, multiple toxin gene clusters were upregulated, including RelE (UniRef90_E7RYC5), RelE/ParE (UniRef90_U2C1G8), RelE/StbE family members (UniRef90_B9D2N3, UniRef90_W2VC95), VapC toxin (UniRef90_UPI00036A705E), an addiction module toxin (UniRef90_F9EDQ2), a putative RNase toxin (UniRef90_A0A318HQ74), and a PIN domain toxin (UniRef90_U7V4D6).

In contrast, Healthy-CI samples displayed upregulation only of toxin-associated transcripts and not antitoxins. These included HicA (UniRef90_A0A0S2ZM78), TelA (UniRef90_A0A133ZXA4), and four PIN family toxins (UniRef90_A0A241Q1Z2, UniRef90_A0A246EGZ2, UniRef90_A0A2B7YY17, UniRef90_A0A2N6TM57), along with an uncharacterized TA system protein (UniRef90_A0A2N8NDD8). No antitoxin gene clusters were differentially upregulated in Healthy-CI, and none of the RHH-domain antitoxins observed in Healthy-CF were present.

While no UniRef90 gene clusters were commonly upregulated across both states, several toxins belonged to shared families, such as PIN domain effectors, suggesting partial convergence at the functional family level despite gene-level exclusivity. These results indicate niche-dependent transcriptional patterns of TA systems among clinically healthy individuals, with broader TA module activation in Healthy-CF and toxin-only enrichment in Healthy-CI.

### Differential Expression of TA Genes Between Healthy and Caries Groups

Across both datasets, transcriptional differences in TA gene cluster expression between caries (Ev: CAa, CAi, CAs, and Dieguez: Caries–Baseline) and healthy (Ev: Healthy–CF, Healthy–CI, and Dieguez: Healthy–Baseline) states were substantial, yet the direction of regulation varied markedly depending on the healthy reference group. In the Ev dataset, caries samples consistently exhibited upregulation of TA genes when compared to the Healthy–CI group (89%), whereas the same caries conditions showed broad transcriptional repression relative to Healthy– CF (average 91.5%). In the Dieguez dataset, when comparing Caries-Baseline vs. Healthy-Baseline (69.33% downregulated), the regulation was modest and followed the trend similar to the Healthy-CF comparisons. This inversion of expression trends was consistent across all spatial caries subtypes (CAa, CAi, CAs), indicating that TA system activity in the caries state is highly contingent on the local reference microbiome (**Table 3**).

There were fewer overlaps of exact gene clusters among the Caries vs. Healthy groups evaluated (***Figure 2B-F***). However, based on the TA families, there are some interesting patterns. PIN domain toxins appear as either upregulated or downregulated depending on contrast, but are consistently differentially expressed across all comparisons, indicating high responsiveness. RHH antitoxins and RelE/StbE family toxins were primarily downregulated in most of the Ev dataset comparisons. These patterns of differential expression reflect that these TA gene clusters may be sensitive to subtle environmental changes and responsive to the broader oral microbiome niche. The Caries-Baseline vs. Healthy-Baseline and all the Caries vs. Healthy-CF contrasts had a consistent pattern with RelE/ParE toxin, RelE toxin, HicB antitoxin being downregulated in these groups. On the other hand, these same families were upregulated in all the Caries vs. Healthy-CI comparisons. This could be a restored memory of activation during caries exposure based on the patient history.

Bacterial toxin RNAse 23 toxin, PHD/YefM antitoxin, VapB antitoxin, YwqK antitoxin and some unannotated antitoxin were differentially expressed only in caries comparisons with Healthy-CI reflecting to a subgroup-specific induction of TA loci, potentially driven by ecological or host-derived pressures unique to the Healthy-CI group. In contrast, no gene cluster was uniquely upregulated in Healthy-CF or Healthy-Baseline contrasts, underscoring the distinctiveness of the CI transcriptional signature.

PHD domain antitoxins were found to be downregulated in Caries-CAa vs. Healthy-CF and Caries-CAs vs. Healthy-CF and all caries subtypes vs. Healthy-CI. They were not significantly expressed in Caries-CAi vs. Healthy-CF, highlighting the environmental change in Caries-CAi. Similar unique gene clusters including Endoribonuclease YoeB (UniRef90_H1LVS8, HMPREF9099_01571), GNAT family toxin (UniRef90_J0MQ72, HMPREF1318_2230), and Fic toxin (UniRef90_L1MBX8, HMPREF9997_02209) were found to be present only in the Caries-CAi vs. Healthy-CF contrast. Interestingly, AbiiEii/AbiGii toxin, Zeta toxin, Holin-like toxin, TelA, Diarrheal toxin, and VapC toxin were found only in the Ev dataset comparisons, and not in the Caries-Baseline vs. Healthy-Baseline comparisons from the Dieguez dataset, reflecting the change could be patient-specific or due to other confounders. In the Dieguez dataset, Baseline comparisons revealed unique downregulation of RelE/ParE toxin (UniRef90_L1NU03, HMPREF9075_02288), RelE toxin (UniRef90_R9RDW5, HMPREF0409_00063), HicA toxin (UniRef90_UPI0007E347B6, KX935_07940), HicB antitoxin (UniRef90_W2CII2, T230_10240), consistent with early suppression of persistence modules in untreated caries.

Notably, Healthy-CI contrasts did not exhibit any uniquely downregulated protein groups, but rather shared suppression with Healthy-CF, reinforcing the earlier finding that TA gene activation, not repression, is more distinctive in Healthy-CI contrasts. While several TA gene clusters exhibited contrast-specific expression, gene-level patterns revealed substantial variability within families, with distinct UniRef90 gene clusters showing divergent regulation even within the same gene cluster. This showcases that TA system dynamics are both family- and gene-specific, shaped by the genomic context and regulatory architecture of individual strains.

### Comparison of Differential Expression among Various Carious Lesion Sites

Spatial comparisons between carious lesion types (CAa, CAi, CAs) revealed widespread upregulation of TA system gene clusters. The strongest upregulation was observed in CAa and CAi when compared to CAs, with over 97% of differentially expressed TA genes upregulated in these contrasts. While the CAa vs. CAi comparison involved fewer differentially expressed genes overall, the directionality of expression favored CAa, underscoring a heightened TA system activation in active lesions. These findings highlight the potential role of TA-mediated stress response pathways in distinguishing active disease states from inactive or healthy sites within the caries continuum (**Table 3**).

A core set of TA protein families was consistently upregulated across all three spatial comparisons including the PIN family toxins, RelE/StbE family toxins, RelE toxins, HicA and HicB toxins, RHH domain antitoxins, PHD domain antitoxins, Xre antitoxins, AbiEii/AbiGii toxins, GNAT family toxins, Holin-like toxins, and YwqK antitoxins, indicating broadly elevated TA gene expression across caries-affected sites relative to sound surfaces.

Several TA gene clusters were also specifically upregulated in individual contrasts. The CAa vs. CAi comparison, showed fewer differentially expressed genes but a clear directional bias toward higher expression in CAa including gene clusters AbrB antitoxin (UniRef90_C8NI31, HMPREF0444_1576), Death-on-curing family protein (UniRef90_UPI00020F0401, PGTDC60_1961), Endoribonuclease YoeB (UniRef90_H1LVS8, HMPREF9099_01571), and YafQ toxin (UniRef90_UPI000479641D). In contrast, CAa vs. CAs had only one unique gene cluster, Bacterial toxin 44 domain-containing protein (UniRef90_S3BAE8, HMPREF1528_02172), while CAi vs. CAs featured Unannotated Antitoxin (UniRef90_A0A1B6W0U8, A7Q00_02305), Diarrheal toxin (UniRef90_F3UU36, yukA), Exfoliative toxin (UniRef90_A0A2X4AK02, shetA), Immunity protein 43 of polymorphic toxin system (UniRef90_A0A2T5XRS3, C8P65_12326), MqsA toxin (UniRef90_F3B3C8, HMPREF0491_01542), Panacea domain-containing protein (UniRef90_UPI000E598125), Unknown Toxin (UniRef90_F2UVA9, HMPREF0059_00360), UDP-N-acetylglucosamine kinase (UniRef90_U7V776, HMPREF0742_00261), VapC toxin (UniRef90_UPI0002B6BC8E, BVwin_10950), and Zeta toxin (UniRef90_C7M709, Coch_0500).

In contrast, downregulated TA gene clusters were rare and observed only in isolated comparisons. The RHH domain antitoxin (UniRef90_U2LXT4, HMPREF1992_01469; UniRef90_U2LXT4, HMPREF1992_01469) was the only family consistently downregulated in both CAa vs. CAi and CAa vs. CAs comparisons. However, these RHH antitoxin genes were found in the upregulated contrasts. Other repressed gene clusters, including Holin-like toxin (UniRef90_V2Y0N7, GCWU0000282_003163) and PHD domain antitoxin (UniRef90_W2VDD7, HMPREF1495_1828) were specific to a single contrast, indicating some specific transcriptional repression of TA systems was limited and localized. These results point toward heightened stress response activation towards active lesion sites (***Figure 2G*)**.

### Treatment-induced Gene Expression Changes in Healthy and Caries Groups

To assess treatment effects on TA-gene expression, we analyzed the longitudinal Dieguez dataset across three time points in both healthy (Healthy-Baseline, Healthy-Fl, Healthy-Fl-Ar) and caries groups (Caries-Baseline, Caries-Fl, Caries-Fl-Ar). Although Dieguez is longitudinal, we performed pairwise contrasts across time points in each group (Healthy-Baseline vs. Healthy-Fl, Healthy-Fl vs. Healthy-Fl-Ar, Healthy-Baseline vs. Healthy-Fl-Ar, Caries-Baseline vs. Caries-Fl, Caries-Fl vs. Caries-Fl-Ar, Caries-baseline vs. Caries-Fl-Ar) and at the same time point between groups (Caries-Baseline vs. Healthy-Baseline, Caries-Fl vs. Healthy-Fl, Caries-Fl-Ar vs. Healthy-Fl-Ar) to keep the analysis comparable to the cross-sectional Ev dataset.

In the Healthy-Baseline vs. Healthy-Fl comparison, 24 TA gene clusters were significantly upregulated, including multiple members of the PIN family toxin (n = 9), RHH-domain antitoxin (n = 6), RelE toxin (n = 3), HicA toxin (n = 3), HicB antitoxin (n = 1), Xre antitoxin (n = 1), and RelE/StbE toxin (n = 1). Only two genes were downregulated in this contrast: RHH domain antitoxin (UniRef90_L1P8R8, HMPREF9073_02960) and Xre antitoxin (UniRef90_L1PPR7, HMPREF9073_01403). In the Healthy-Baseline vs. Healthy-Fl-Ar comparison, the directionality reversed, with 14 TA genes downregulated and only two upregulated RHH-domain antitoxins (UniRef90_U1R3T5, HMPREF9065_00394; UniRef90_G6C504, HMPREF9093_01650). The downregulated genes in this contrast included multiple RHH-domain antitoxins (n = 5), as well as HicA toxin (UniRef90_D7NFI9, HMPREF0665_02330), HicB antitoxin (UniRef90_L1PKY8, HMPREF9073_01869), PHD domain antitoxin (UniRef90_L1PUB1, HMPREF9073_00941), PIN family toxin (UniRef90_A0A2C6BMA4, CBG54_02990), RelE toxin (UniRef90_A0A096ASI9, HMPREF0661_05145), ToxN toxin (UniRef90_E8KF34, HMPREF0027_0451), Xre antitoxin (UniRef90_U1PUH2, HMPREF0043_00600) and two Holin-like toxins (UniRef90_A0A134A313, HMPREF3186_00500; UniRef90_S2ZQS1, HMPREF1477_01317). A similar pattern was observed in the Healthy-Fl vs. Healthy-Fl-Ar contrast, where 10 TA genes were downregulated and only three were upregulated: Bacterial toxin 44 domain-containing protein (UniRef90_E1KT92, HMPREF9296_0290), RelE/StbE family toxin (UniRef90_U2R2D9, HMPREF1552_02277), and RHH domain antitoxin (UniRef90_G6C504, HMPREF9093_01650).

Importantly, several TA gene families exhibited directional transitions across timepoints. Members of the PIN family toxin, RelE toxin, and HicA toxin families were consistently upregulated in the Baseline vs. Fl contrast but downregulated in later comparisons. RHH-domain antitoxins were the most frequently modulated, with eight upregulated and seven downregulated instances across comparisons, involving distinct UniRef90 gene identifiers. RHH domain antitoxins were consistent with the pattern of sensitive change to slight environmental variation as observed in Healthy vs Caries comparisons. A small number of genes showed stable directionality across timepoints. Several TA-associated gene clusters were differentially expressed in more than one contrast. PIN family toxin (UniRef90_A0A2G9FAK1, CI114_08815) and HicA toxin (UniRef90_A0A2C6BV61, CA836_05975) were both upregulated in Baseline vs. Fl and downregulated in Fl vs. Fl-Ar. Other RHH-domain antitoxins that were differentially expressed across treatment stages included UniRef90_U1R3T5 (HMPREF9065_00394) and UniRef90_G6C504 (HMPREF9093_01650), both upregulated in Baseline vs. Fl and again in Baseline vs. Fl-Ar. UniRef90_L1P8R8 (HMPREF9073_02960) and UniRef90_L1NZV5 (HMPREF9078_00729) both downregulated in Baseline vs. Fl-Ar and Fl vs. Fl-Ar (**Figure 2H)**. The number of upregulated TA gene cluisters declined markedly across the timepoints (24 in Baseline vs. Fl, 3 in Fl vs. Fl-Ar, 2 in Baseline vs. Fl-Ar), while downregulated gene clusters increased (2, 10, and 14, respectively), reflecting a shift from broad early induction to widespread suppression. Antitoxin gene clusters were more frequently differentially expressed than toxin gene clusters across all stages, suggesting that TA gene modulation in healthy individuals is dominated by antitoxin-associated regulation during and after exposure to fluoride and arginine.

In the Caries group, TA gene expression changed markedly following Fl and Fl-Ar treatments. In the Caries-Baseline vs. Caries-Fl comparison, a total of 10 genes were significantly differentially expressed, including 4 downregulated and 6 upregulated TA gene clusters. The downregulated gene clusters included PIN family toxin (UniRef90_A0A2B7YNI3, CI114_10250), RHH domain antitoxin (UniRef90_U1R9S4, HMPREF0043_01920), HicA toxin (UniRef90_U2PKD9, HMPREF1552_01229), and a HicB antitoxin (UniRef90_W2CII2, T230_10240). The upregulated gene clusters comprised of three PIN family toxins (UniRef90_A0A2C5ZYQ6, CBG52_10650; UniRef90_A0A2C6C3I1, CBG56_11515; UniRef90_A0A2N6TM57, CJ209_03975), one RelB/DinJ antitoxin (UniRef90_A5TWG4, FNP_1457), one RHH domain antitoxin (UniRef90_J6IUG2, HMPREF1154_2041), and one RelE/StbE family toxin (UniRef90_X8HWZ7, HMPREF1501_0456). In the Caries-Baseline vs. Caries-Fl-Ar contrast, 13 TA gene clusters were upregulated and 3 were downregulated. Upregulated gene clusters included PIN family toxins (n=4), RelE/StbE family toxins (n = 2), RHH-domain antitoxin (n = 2), and one each of RelE toxin, RelB/DinJ antitoxin, and PHD domain antitoxin. Downregulated gene clusters included RelE/ParE toxin (UniRef90_L1NU03, HMPREF9075_02288), RelE toxin (UniRef90_U1SAA0, HMPREF9065_01198), and HicA toxin (UniRef90_U2PKD9, HMPREF1552_01229). The Caries-Fl vs. Caries-Fl-Ar comparison showed 5 TA gene clusters upregulated and 2 downregulated. Upregulated gene clusters were two PIN family toxins (UniRef90_A5TV72, FNP_1001; UniRef90_E6LLD9, HMPREF0381_0774), a RelE/StbE family toxin (UniRef90_D3QZD1, HMPREF0868_1572), an RHH domain antitoxin (UniRef90_G9ZCM6, HMPREF9080_00502), and a PHD domain antitoxin (UniRef90_W2VDD7, HMPREF1495_1828)). Downregulated gene clusters were a PIN family toxin (UniRef90_A0A2C6C3I1, CBG56_11515) and an RHH domain antitoxin (UniRef90_J6HZ39, HMPREF1154_0792).

Across all three caries comparisons, TA gene regulation exhibited family-specific directional trends. The PIN family toxins were the most frequently regulated, with 6 gene clusters upregulated across all comparisons and 2 downregulated, indicating a largely inducible profile. The only shift in the direction was observed in PIN family toxin (UniRef90_A0A2C6C3I1, CBG56_11515), which was upregulated in Baseline vs. Fl and was downregulated in Fl vs. Fl-Ar. RelE/StbE family toxins were consistently upregulated, with no instances of downregulation, appearing in all three contrasts (UniRef90_D3QZD1, HMPREF0868_1572; UniRef90_X8HWZ7, HMPREF1501_0456). Conversely, HicA toxin was consistently downregulated in both Baseline vs. Fl and Baseline vs. Fl-Ar (UniRef90_U2PKD9, HMPREF1552_01229). RelE toxins showed mixed regulation: upregulated in Baseline vs. Fl-Ar (UniRef90_A5TVF0, FNP_1085) and downregulated in the same contrast (UniRef90_U1SAA0, HMPREF9065_01198), suggesting possible gene-specific divergence in response. RHH-domain antitoxins were both up- and downregulated across comparisons, with gene-level specificity. PHD domain antitoxin (UniRef90_W2VDD7, HMPREF1495_1828) was upregulated in both Baseline vs. Fl-Ar and Fl vs. Fl-Ar (***Figure 2I*)**.

The trend across the treatment timeline (Baseline to Fl to Fl-Ar) in caries samples showed predominant upregulation of TA genes, in contrast to healthy individuals, where suppression dominated later stages. Several TA genes appeared in multiple contrasts and maintained directionality. This suggests that, in the caries microbiome, certain TA systems remained transcriptionally active throughout treatment exposure, with little evidence of resolution or suppression observed in the healthy group.

Healthy individuals exhibited a broad early induction of TA systems in response to fluoride, predominantly involving antitoxins and translational toxins (RelE, PIN, HicA). In caries samples, the transcriptional response was more mixed, with HicA and HicB families downregulated, while RelB/DinJ and RelE/StbE families were upregulated, suggesting disease-modulated responsiveness or baseline dysregulation in the caries microbiome (***Figure 2H-I*, Green circles)**. This contrast revealed an inverse transcriptional shift between groups. Healthy samples showed widespread TA gene suppression by the Fl-Ar stage, suggesting transcriptional resolution or stress adaptation. In contrast, caries samples maintained or amplified TA activation, with continued upregulation of PIN, RelE, RelE/StbE, and RHH-domain antitoxins, indicating a sustained or non-resolving transcriptional activation even at the late treatment stage (***Figure 2H-I*, Orange circles)**. While healthy samples showed a predominantly suppressive response between Fl and Fl-Ar stages, caries samples maintained ongoing TA activation, especially of RelE/StbE, PIN, PHD, and RHH-domain antitoxins. Only two genes were suppressed in caries, suggesting the absence of resolution dynamics seen in healthy samples and instead a persistent activation trajectory (***Figure 2H-I*, Blue circles)**.

### Comparative Treatment-induced Gene Expression Changes in Healthy vs. Caries Groups

To assess the changes longitudinally, we evaluated the changes at each time point in the caries and healthy groups (Caries-Baseline vs. Healthy-Baseline, Caries-Fl vs. Healthy-Fl, Caries-Fl-Ar vs. Healthy-Fl-Ar). The baseline stage, as explained at the Healthy vs. Caries comparisons, suggests a pre-treatment imbalance in caries samples, marked by elevated expression of translational toxins (HicA, PIN) and suppression of regulatory antitoxins, especially from the RHH and RelE families.

At the fluoride stage, the transcriptional contrast widened, with caries samples showing only two upregulated gene clusters, all PIN family toxins, and a bigger chunk of 12 downregulated TA gene clusters. The downregulated set encompassed a broad suite of RHH-domain antitoxins (n = 6), Xre antitoxins (n = 2), and one each of RelE toxin, RelE/StbE toxin, AbiEii/AbiGii toxin, and an additional HicB antitoxin. This shift reflects a disproportionate suppression of transcriptional repressors and canonical antitoxins in caries under fluoride exposure in contrast to the healthy group, where these gene clusters remain stably or moderately expressed. Specifically, no RelE, HicA toxins, or RHH-domain antitoxins were upregulated in caries at this stage, indicating a flattened transcriptional stress response despite chemical perturbation.

In the Fl–Ar stage, the caries group exhibited a rebound activation, with 11 TA gene clusters upregulated and 5 downregulated relative to healthy samples. Upregulated gene clusters included multiple HicA toxins (UniRef90_A0A0S2ZM78, RN87_05845; UniRef90_A0A0X8JSY8, AXF11_00955; UniRef90_A0A2N6TLH8, CJ209_02680), RelE toxins (UniRef90_A0A250FNZ8, CGC50_06225; UniRef90_U1SAA0, HMPREF9065_01198), RHH domain antitoxins (UniRef90_D3IKU4, HMPREF0670_02113; UniRef90_J6HZ39, HMPREF1154_0792), YwqK antitoxins (UniRef90_A0A241Q2B3, CBG61_08500; UniRef90_A5TSM3, FNP_0079), a PIN family toxin (UniRef90_A0A2N6TM57, CJ209_03975), and a Bacterial toxin 44 domain-containing protein (UniRef90_E1KT92, HMPREF9296_0290). Downregulated gene clusters included HicA toxin (UniRef90_UPI0007E347B6, KX935_07940), a HicB antitoxin (UniRef90_U1Q2Z8, HMPREF0043_01620), a PIN family toxin (UniRef90_E6LLD9, HMPREF0381_0774), an RHH domain antitoxin (UniRef90_A0A0M3UUY0, CI114_09045), and Unknown Toxin-antitoxin system protein (UniRef90_W2CTG9, T230_04075), suggesting continued but partial suppression of canonical regulators. Interestingly, several HicA toxins showed contradictory expression across stages, upregulated at baseline but downregulated at Fl–Ar, indicating dynamic modulation across treatments **(*Figure 2J***, **Table 3**.**Table 3, Supplementary tables S2, Supplementary figures S1, S3)**.

Overall patterns revealed a consistent overexpression of HicA and PIN family toxins in caries relative to healthy samples across all treatment stages, highlighting their potential role in caries-associated stress adaptation. In contrast, RHH-domain and Xre-type antitoxins were consistently downregulated in caries, suggesting a loss of repression machinery. While healthy samples exhibit homeostatic or suppressive regulation of TA systems following treatment (as observed in prior within-group comparisons), caries samples appear to mount a delayed and dysregulated activation, with elevated toxin expression emerging only at the Fl–Ar stage. This trajectory suggests that caries-associated microbiomes exhibit disrupted TA regulatory balance, particularly within PIN–RHH, HicA–HicB, and RelE–RelB/DinJ modules. Detailed visualizations and tables are presented in the supplementary figures (**S1-S4**) and supplementary tables **(S2-S4).**

### Functional Insights into TA Gene Clusters

To assess the functional relevance of the differentially expressed TA gene clusters, we cross-referenced them with TADB3, a curated repository of experimentally validated and computationally predicted TA genes and proteins. For each significant TA gene cluster, we evaluated its overlap with TADB3 entries and recorded the nature of the supporting evidence (experimental vs. computational). Using DIAMOND BLASTp, we queried the TADB3 reference set against the UniRef90 clusters detected in our metatranscriptomic data analysis, applying a stringent filter of ≥90% sequence identity and selection of the lowest e-value per match. This yielded 18 unique UniRef90 gene clusters with validated TA system annotations, comprising 11 toxins and 7 antitoxins. All validated genes were significantly differentially expressed in at least one contrast in the Dieguez or Ev datasets **(*Figure 3***, ***Figure 4***; **Table 3**, ***Table 4*)**. In these validated genes clusters, HicA, RelE, YafQ, VapC toxins and HicB, RelB antitoxins were detected in both Dieguez and Ev datasets, indicating shared TA-associated responses across different populations and study designs.

**Figure 3.**
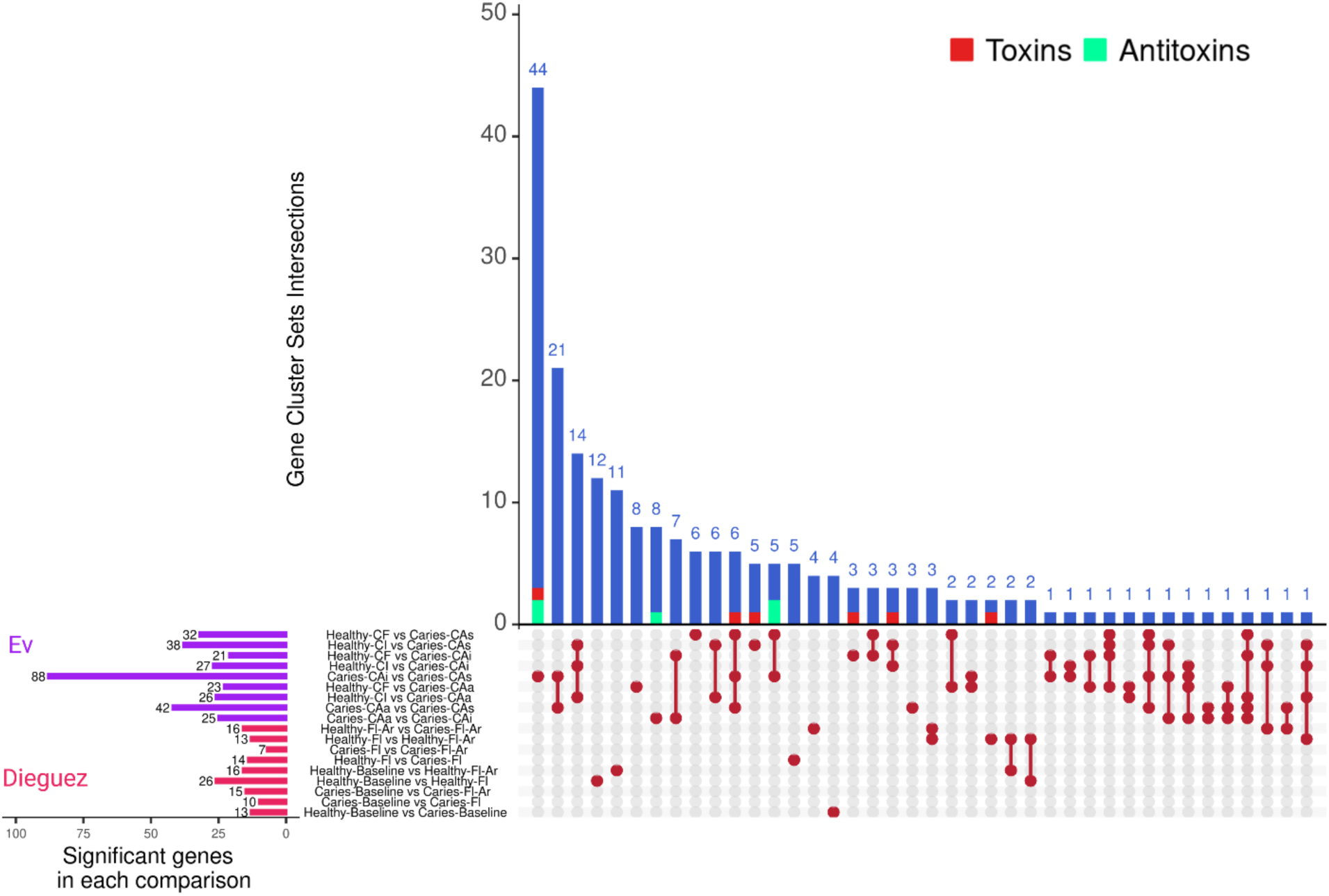
Shared and unique differentially expressed TA system gene clusters across condition comparisons in Dieguez and Ev datasets. UpSet plot showing the intersection of significant TA system gene clusters across condition comparisons in Ev (purple) and Dieguez (pink) datasets. Horizontal bars (left) represent the total number of differentially expressed gene clusters in each contrast. Vertical bars (top) indicate the number of overlapping differentially expressed gene clusters among different condition comparisons. Red and green bars highlight genes validated as toxins or antitoxins in the TADB3 database, respectively.

**Figure 4.**
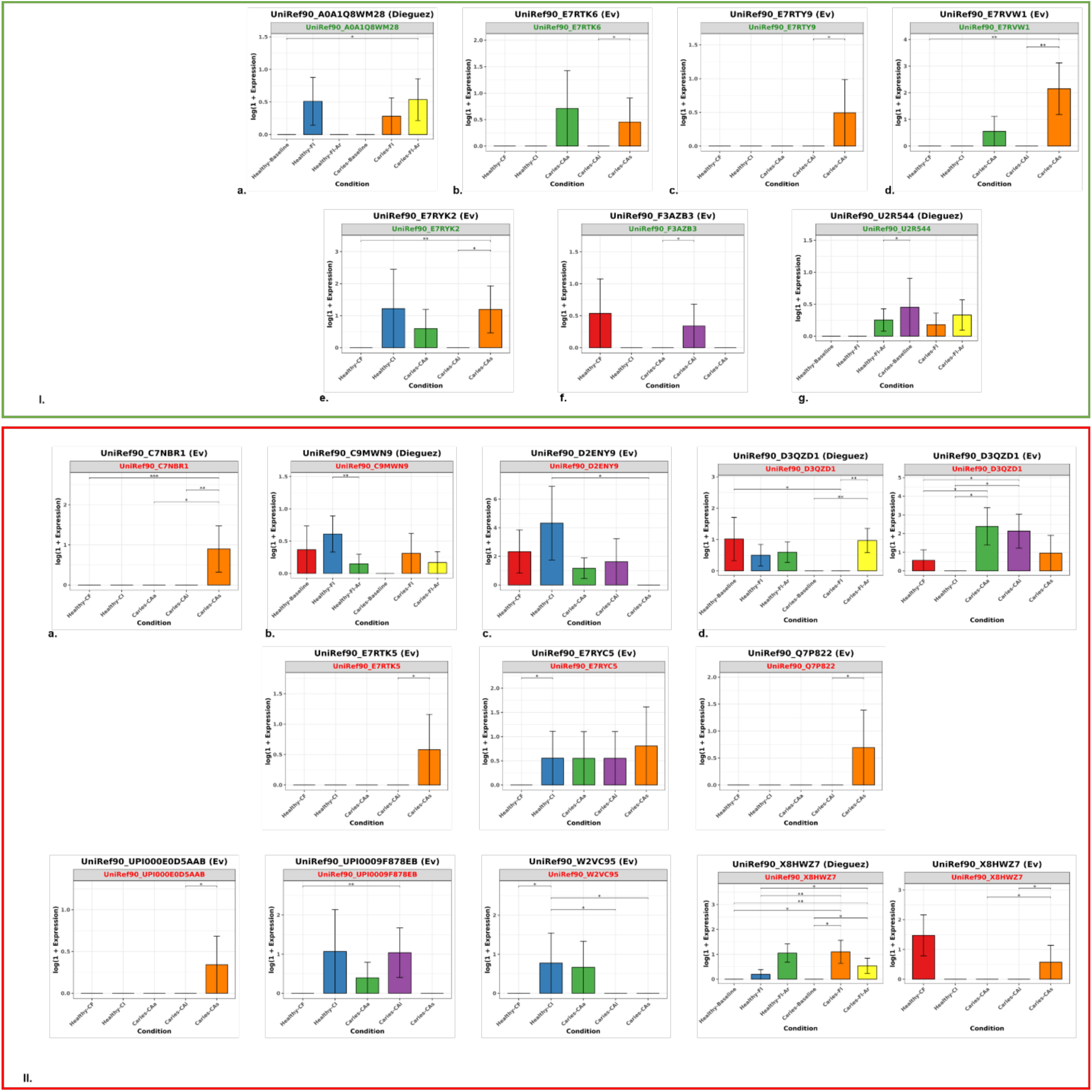
Differential expressions of validated toxin and antitoxin genes across conditions in Dieguez and Ev datasets. Barplots displaying log-transformed normalized expression values (log1p (log(1+x))) for experimentally validated or computationally predicted toxins and antitoxins listed in ***Table 4***. Genes in green box (I, panels a-g) correspond to antitoxins, while those in red box (II, panels a-k) correspond to toxins. Each panel shows expression profiles across condition-specific groups in either Dieguez or Ev dataset. Error bars represent variability across samples. Black brackets with significance stars (p < 0.05, p < 0.01, p < 0.001) denote statistically significant differences between conditions based on ANCOM-BC analysis. X-axis labels reflect dataset-specific condition orders to ensure consistency in comparative interpretation.

**Table 4.**
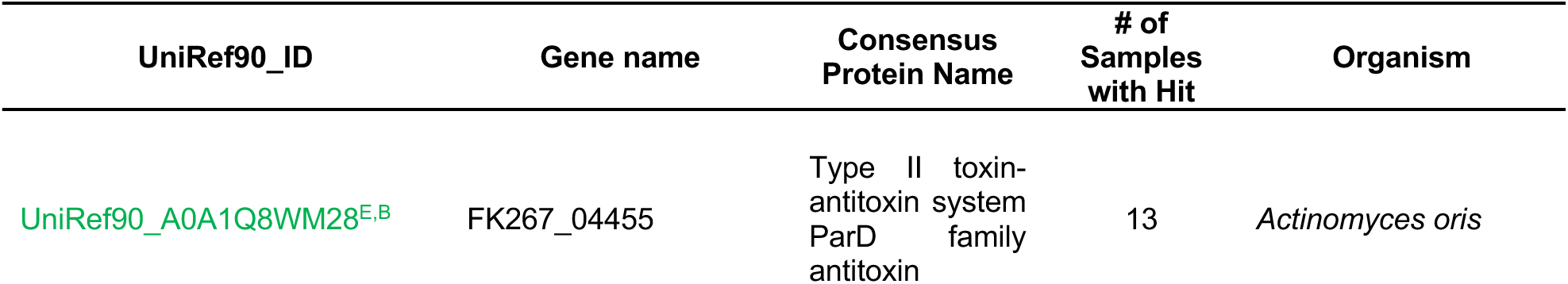
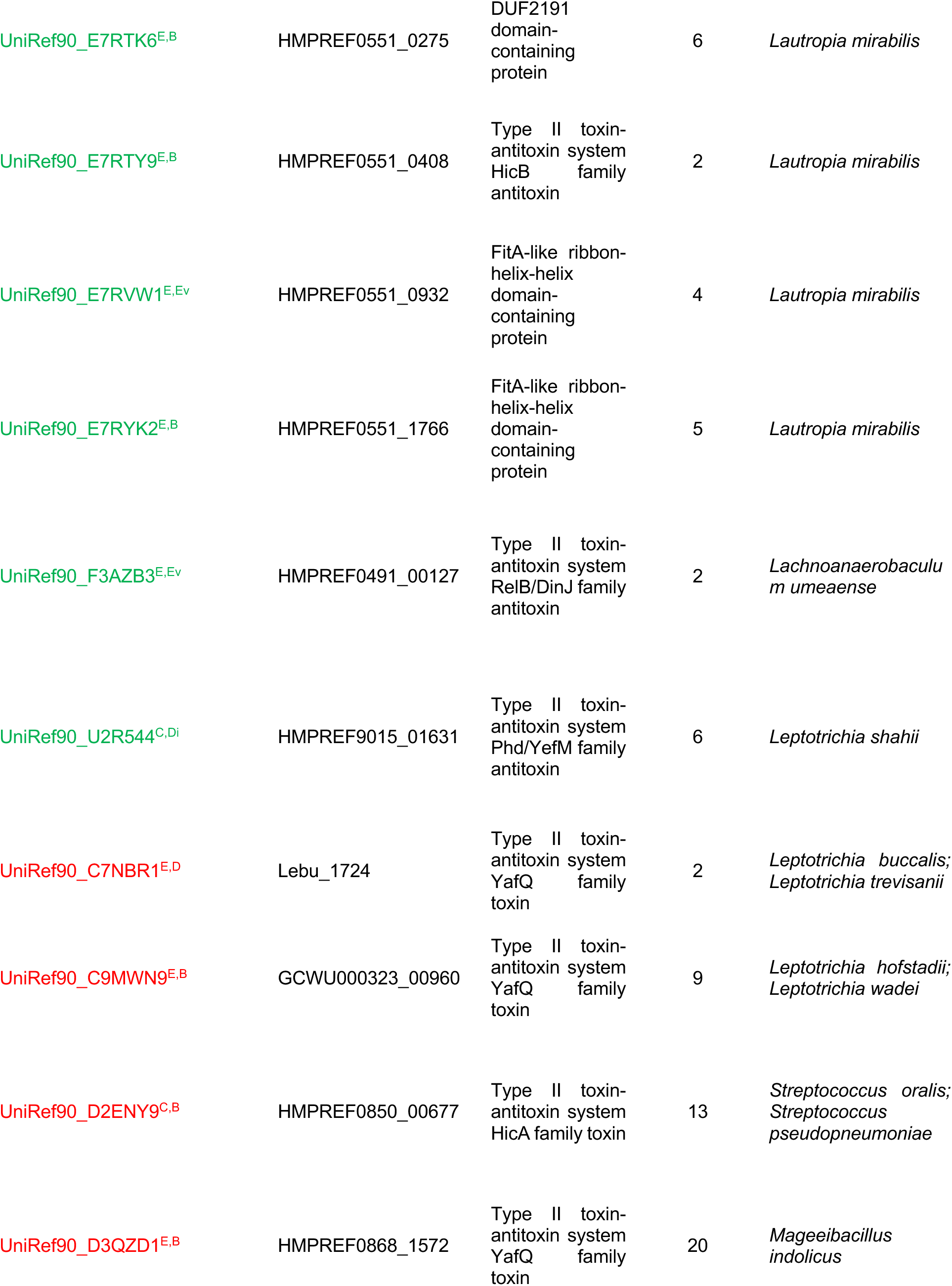
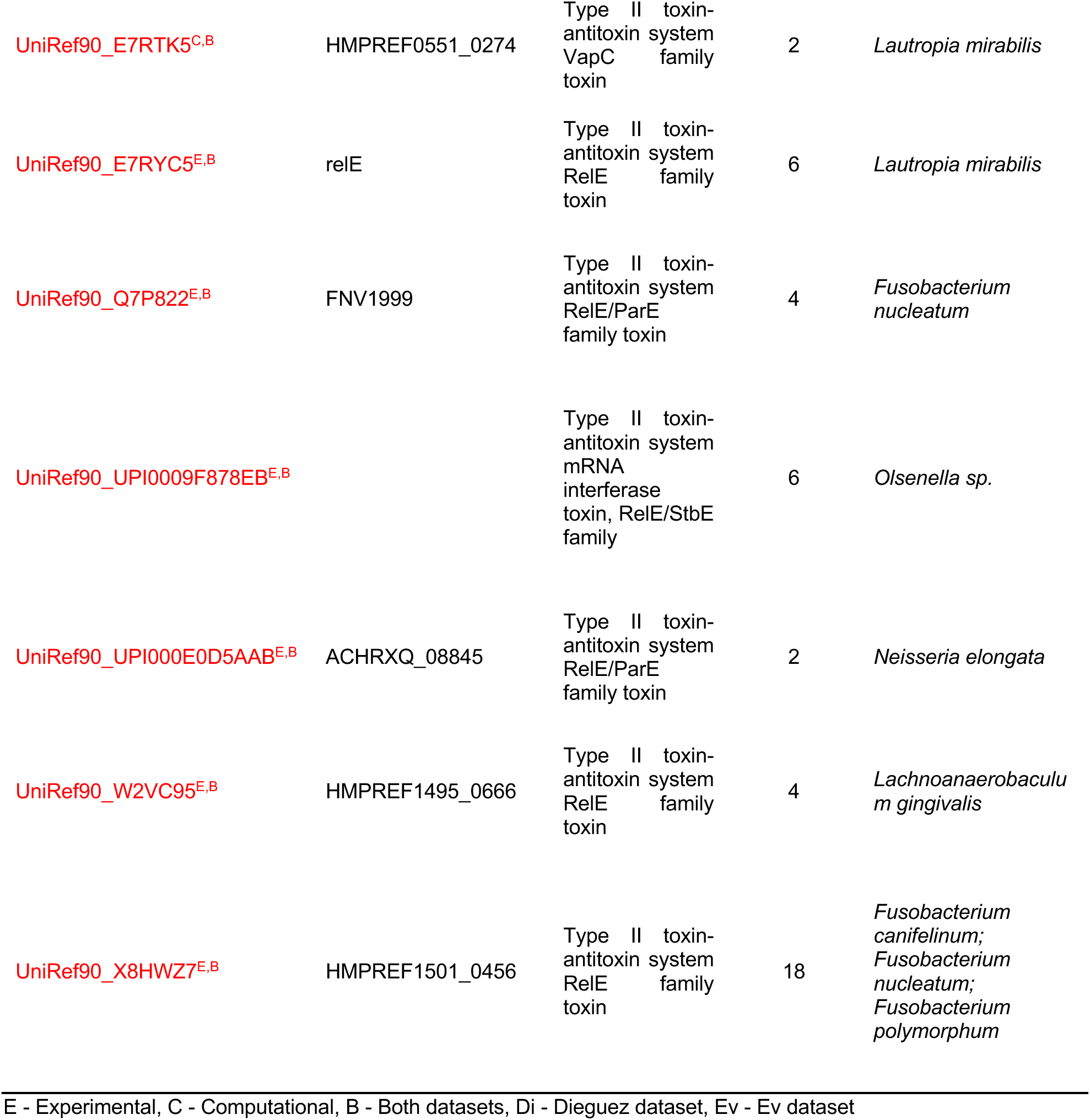
Experimentally or computationally supported TA genes detected in the oral microbiome and validated through TADB3. High-confidence TA system gene clusters were identified by aligning differentially expressed UniRef90 sequences against the TADB3 database using DIAMOND. Complete annotation details, including accessions and alignment scores, are provided in supplementary tables S3 and S4.

Condition-specific analyses revealed that several of these validated gene clusters were associated with disease-state transitions in Ev dataset. The comparison between Caries-CAi and Caries-CAs showed the largest number of significant TA gene cluster changes, including 88 differentially expressed clusters, of which six matched validated TADB3 entries such as HicA and HicB. Additional comparisons, including Caries-CAa versus -CAi and -CAa versus -CAs, revealed 25 and 42 differentially expressed TA clusters, respectively. Two to three validated toxins were identified per contrast, including members of the YafQ, VapC, and RelE families. Notably, UniRef90_A0A2C5ZYQ6 (ParD antitoxin) and UniRef90_A0A2C6C3I1 (YafQ toxin) were consistently detected in all three binary comparisons between Healthy-CF and Caries-CAa, -CAi, and -CAs, indicating reproducible TA system signatures associated with caries-active states.

In the Dieguez dataset, which captures temporal changes following fluoride treatments, validated TA gene clusters also showed distinct expression patterns. UniRef90_Q7P822 and UniRef90_UPI000E0D5AAB, both RelE/ParE toxin family members, were differentially expressed between baseline and fluoride-treated caries samples. In the healthy arm, UniRef90_F3AZB3 (RelB antitoxin) and UniRef90_A0A1Q8WM28 (ParD antitoxin) showed upregulation following Fl-Ar treatment relative to baseline. Furthermore, UniRef90_D3QZD1 (YafQ) and UniRef90_X8HWZ7 (RelE) were differentially expressed between caries-affected individuals post-Fl and Fl-Ar treatment, suggesting a possible continued role in stress adaptation following intervention. The differential expressions of these validated genes are represented in ***Figure 4***.

These results demonstrate that several differentially expressed TA system gene clusters identified across healthy, diseased, and treatment conditions correspond to experimentally and computationally validated entries in TADB3. The recurrence of these gene clusters across datasets, contrasts, and treatment stages suggests their potential involvement in bacterial stress regulation and adaptive dynamics of the oral microbiome in response to caries progression and therapeutic interventions.

### Expression Changes of the Validated TA system Gene Clusters

Across the 18 validated toxin–antitoxin system genes (11 toxins, 7 antitoxins), consistent differences in expression were observed between caries and healthy conditions across both datasets and treatment stages. In the Dieguez dataset, the toxin UniRef90_D3QZD1 (HMPREF0868_1572, YafQ family toxin, *Mageeibacillus indolicus*) and UniRef90_X8HWZ7 (HMPREF1501_0456, RelE family toxin*, Fusobacterium canifelinum / nucleatum / polymorphum*) were significantly upregulated in caries samples at Baseline, Fl, and Fl–Ar stages, with UniRef90_D2ENY9 (HMPREF0850_00677, HicA family toxin, Streptococcus oralis / pseudopneumoniae) showing the largest increase in Caries–CAs compared to Healthy–CI. Toxins UniRef90_C9MWN9 (GCWU000323_00960, YafQ family toxin, *Leptotrichia hofstadii* / *wadei*) and UniRef90_U2R544 (HMPREF9015_01631, Phd/YefM family antitoxin, *Leptotrichia shahii*) also showed differential expression across treated conditions. Among antitoxins, UniRef90_E7RYC5 (relE, RelE family toxin, *Lautropia mirabilis*) was moderately elevated in Healthy–CI. In the EV dataset, UniRef90_C7NBR1 (Lebu_1724, YafQ family toxin, *Leptotrichia buccalis / trevisanii*), UniRef90_Q7P822 (FNV1999, RelE/ParE family toxin, *Fusobacterium nucleatum*), and UniRef90_E7RTK5 (HMPREF0551_0274, VapC family toxin, *Lautropia mirabilis*) were downregulated in Healthy–CF relative to later caries stages. In contrast, UniRef90_W2VC95 (HMPREF1495_0666, RelE family toxin, *Lachnoanaerobaculum gingivalis*) was consistently upregulated in Caries–CAi and CAs compared to Healthy–CI. Several antitoxins in EV, including UniRef90_E7RTY9 (HMPREF0551_0408, HicB family antitoxin), UniRef90_E7RVW1 (HMPREF0551_0932, FitA-like antitoxin), and UniRef90_E7RTK6 (HMPREF0551_0275, DUF2191 domain antitoxin) from Lautropia mirabilis, were consistently downregulated in caries stages. Notably, UniRef90_D3QZD1 and UniRef90_X8HWZ7 (RelE/StbE family toxin) were differentially expressed in both datasets across multiple comparisons, indicating dataset-consistent transcriptional shifts in these TA systems between Healthy and Caries groups (***Figure 4***, Supplementary tables S2-S4).

### Functional Insights into Significant TA UniRef90 Clusters

To characterize the validated TA gene clusters functionally, we annotated the TADB3 experimentally validated or computationally predicted 18 UniRef90 sequences using InterProScan and eggNOG-mapper **(*Table 5*A-B)**. InterProScan was used to identify conserved protein domains, superfamily assignments, and associated Gene Ontology (GO) terms, while eggNOG-mapper provided orthology-based annotation, COG category classification, and pathway mapping via KEGG BRITE [33]. To comprehensively capture domains, motifs, and pathways, we ran both InterProScan and eggNOG-mapper. Detailed information with the accessions, e-values, and annotations specific to each tool is presented in supplementary table ***S5***.

**Table 5.**
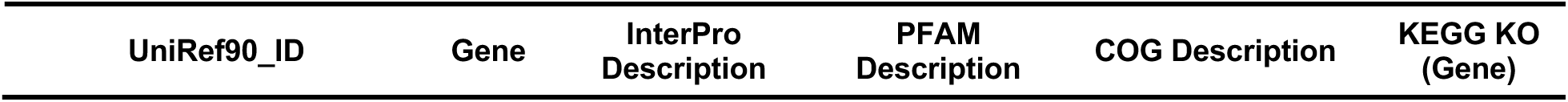
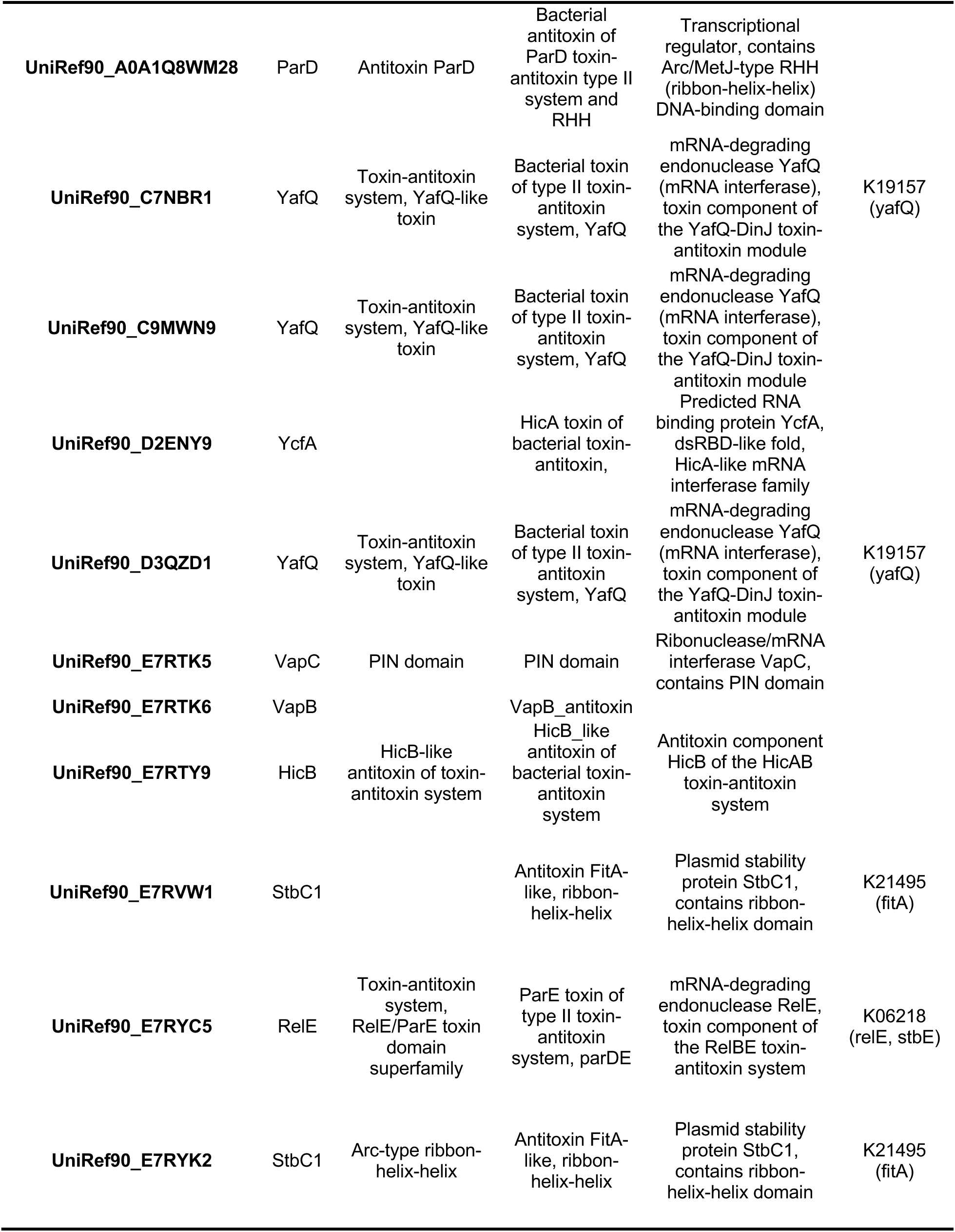
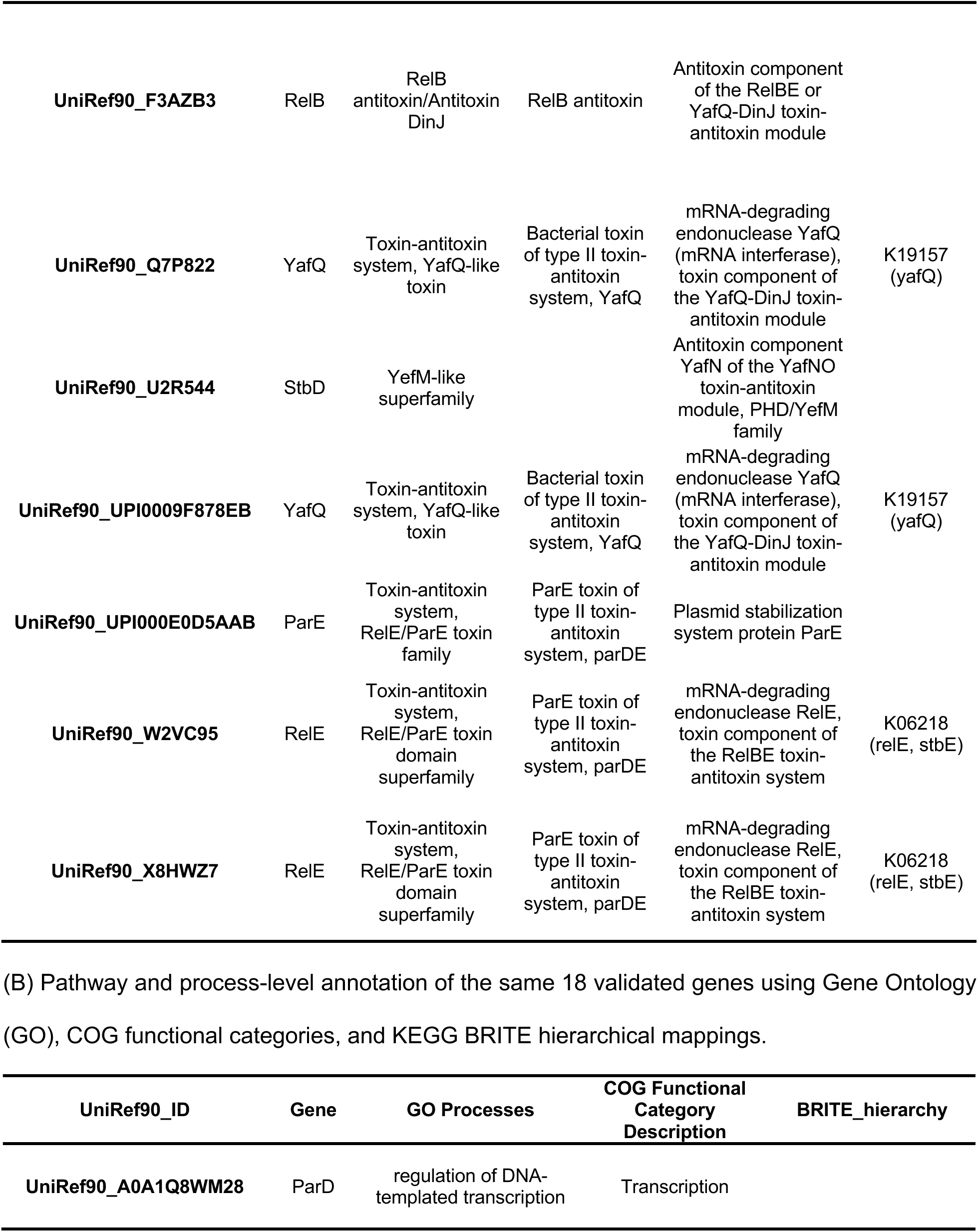
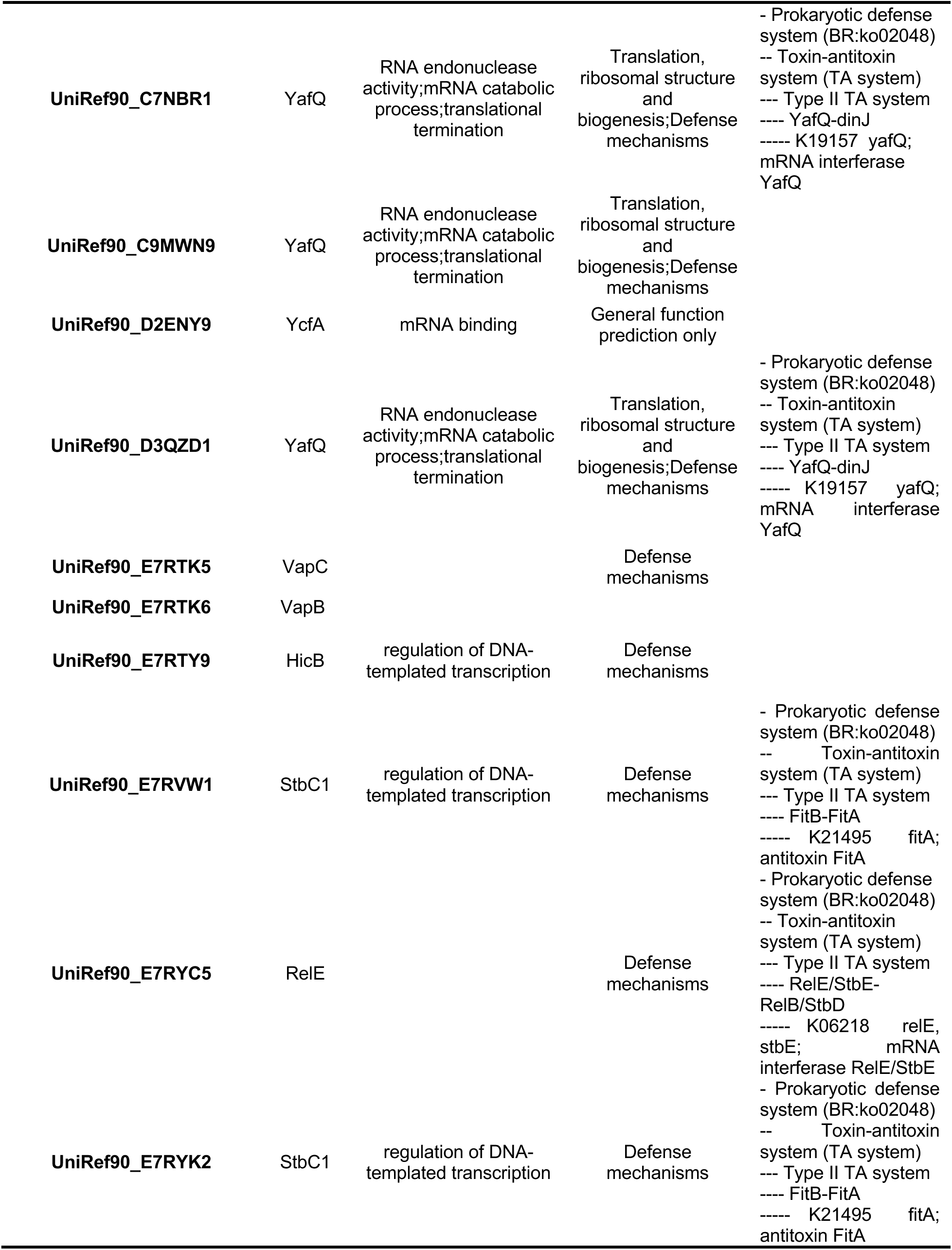
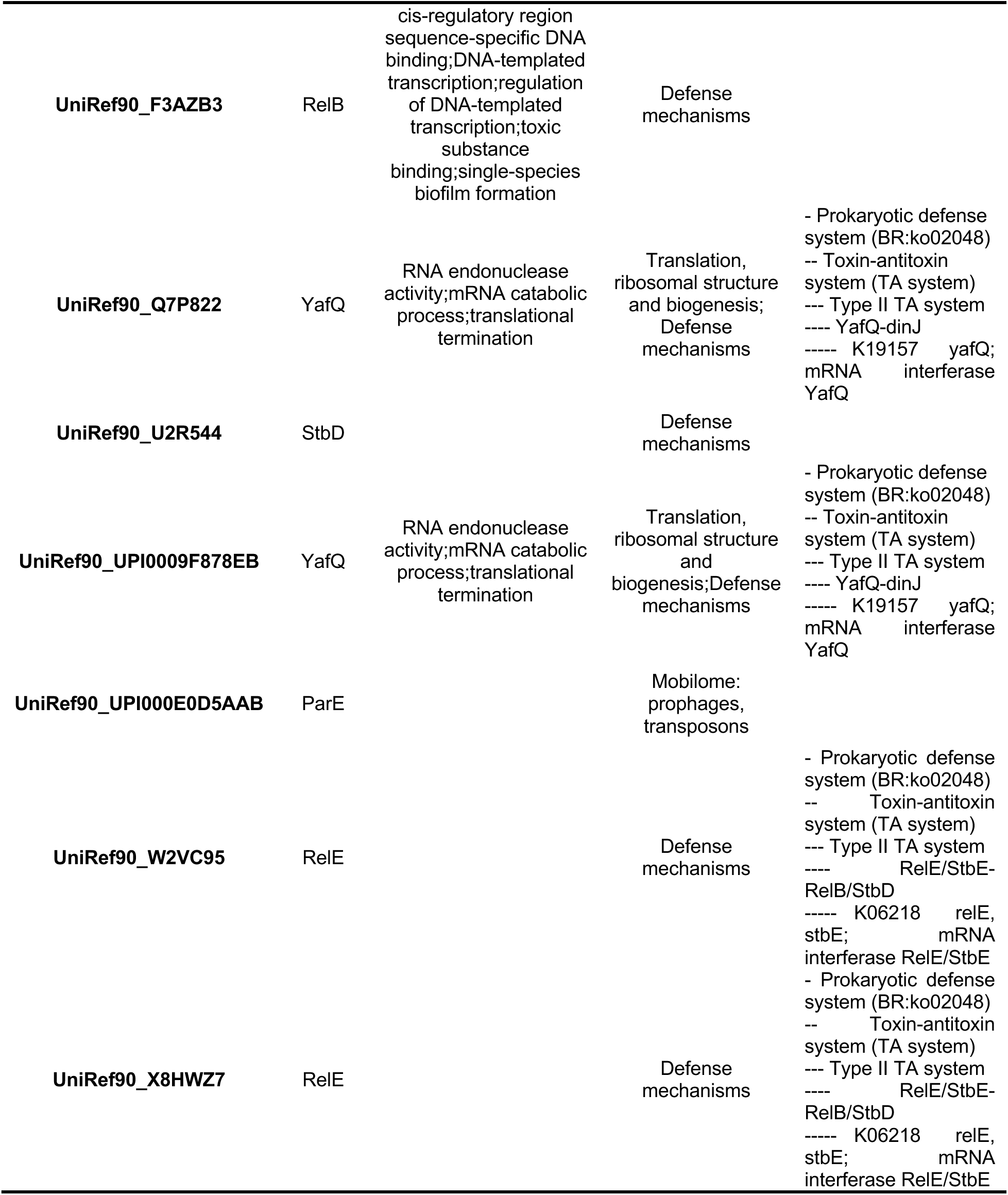
Functional annotation of the shortlisted TA system gene clusters. (A) Summary of domain-level annotations for the shortlisted TA gene clusters using InterProScan and eggNOG-mapper. Each UniRef90 cluster was mapped to consensus protein names and annotated with conserved domains where available.

All eleven toxins were assigned to well-characterized TA superfamilies. Most were annotated as RelE/YafQ/ParE toxin family members, with conserved RelE-like domains and GO terms linked to RNA endonuclease activity, mRNA catabolism, and translational inhibition. These were further classified under COG3041, COG2026, or COG3668, supporting their function as mRNA interferases. Toxins such as HicA and VapC showed distinct domain architectures, including PIN and YcfA-like folds, consistent with ribonuclease activity under stress conditions. KEGG ortholog assignments grouped these toxins into multiple type II TA modules, notably RelE/StbE-RelB/StbD, YafQ-DinJ, and FitB-FitA systems, all nested under the prokaryotic defense hierarchy (ko02048).

Among the seven antitoxins, domain architectures were characterized by ribbon-helix-helix (RHH) motifs, RelB/DinJ-like folds, or YefM superfamilies, aligning with their predicted roles in transcriptional repression. GO terms across antitoxins commonly included regulation of DNA-templated transcription and toxic substance binding, indicating transcriptional regulation of cognate toxins and roles in broader stress response. Antitoxins such as ParD, HicB, and RelB were consistently mapped to COG3609, COG1598, or COG3077, confirming their canonical roles within type II TA systems.

These dual-layered annotations reaffirm that the differentially expressed, validated TA genes detected in our datasets represent functionally active components of known prokaryotic TA systems. InterProScan and eggNOG-mapper provided complementary evidence of domain-specific molecular activity and evolutionary and functional classification. This provides solid evidence to thefunctional context for the observed transcriptional changes.

## Discussion

Our metatranscriptomic analysis revealed that TA systems are active in the oral microbiome and differ between healthy, caries, and treatment conditions. Among these, Type II TA systems (protein toxins with protein antitoxins) are highly prevalent in bacteria and have been extensively studied [34, 35]. Their proteinaceous components often exist in multiple chromosomal copies, likely aiding transcript detection even at modest sequencing depth. Although we did not target Type II TA genes explicitly, they emerged prominently in the data. This underscores the importance of TA modules, known for their roles in stress survival, biofilm formation, and persistence, in the adaptive response of the oral microbiome.

Our findings indicate that oral bacteria modulate TA system expression in response to even subtle environmental changes (comparing healthy, caries-active, and post-treatment states). TA genes identified in our dataset belonged to classic TA families, suggesting these microbes employ dormancy and toxin-antitoxin balancing acts as a strategy to endure hostile conditions. This aligns with prior evidence that core oral species such as *Streptococcus mutans [*17, 18, 36], *Fusobacterium nucleatum [*37, 38], and *Treponema denticola [*12] harbor multiple TA systems linked to stress tolerance and persistence. Our results confirm the presence of such TA genes in oral biofilms and reveal significant expression changes across conditions, indicating TA systems may be crucial factors altering microbial community states during dental caries progression.

This study builds on and extends previous oral microbiome research. Carda-Diéguez *et al.* analyzed plaque before and after Fl or Fl-Ar dentifrice use. They observed minimal changes in overall community diversity at the RNA (metatranscriptome) level, though DNA-based richness indices slightly decreased after treatment. Their interpretation was that bacteria which adapted to the new environment (especially arginine-rich conditions) became dominant, reducing species richness. TA systems provide a mechanistic basis for such adaptability by enabling a subset of cells to enter a stress-tolerant state, TA modules could allow certain bacteria to outcompete others under fluoride or arginine exposure, leading to a less diverse but more resilient community [20]. Similarly, Ev et. al., claimed that caries-free biofilms are functionally enriched in two-component regulatory systems. Two-component systems are sensors for environmental change [21], and our findings suggest that TA systems might be downstream effectors in the same context, helping bacteria survive those sensed changes. Together with prior studies, our findings highlight toxin-antitoxin systems as underappreciated drivers of transitions between healthy, diseased, and treated oral states.

Using the TADB 3.0 database for annotation, we identified a set of putative TA loci in our samples, primarily from the genera *Leptotrichia* (5 loci), *Fusobacterium* (3 loci), *Streptococcus* (2 loci), and *Lachnoanerobaculum* (2 loci). With the exception of *Streptococcus*, these are generally low-abundance taxa that have been observed in either caries-active or healthy oral cavities. The enrichment of TA genes among low-abundance organisms suggests that stress-response modules are utilized even by minor members of the community. *Leptotrichia* spp., are opportunistic anaerobes that thrive in carious environments; their surface protrusions and LPS promote virulence and adhesion, and they tend to increase when urease-producing commensals decline [39–41]. The presence of multiple TA genes in *Leptotrichia* supports the idea that this genus endures the acidic, competitive caries niche by inducing persistence mechanisms. In contrast, *Streptococcus oralis*, a commensal important for initial biofilm formation, is typically abundant in health [20, 42, 43]. TA modules detected in *S. oralis* (and in *S. pseudopneumoniae*, a close relative of the pathogen *S. pneumoniae* [44]) implies that even commensals retain latent capabilities to withstand abrupt stresses such as sugar surges or pH drops.

Interestingly, we also detected TA genes from *Actinomyces oris [*45, 46] and *Neisseria elongata [*40]. Both species have been reported in caries-free and caries-active plaques, indicating context-dependent roles [20, 40, 41, 43, 47]. The presence of TA loci in both healthy and diseased groups suggests a contribution to community resilience independent of caries status. Notably, *Lautropia mirabilis*, a low-abundance bacterium consistently associated with healthy and caries-free children [20, 41], expressed TA transcripts that may underlie its ability to persist as a stable, minor biofilm member despite episodic perturbations. We also observed TA genes from *Fusobacterium* spp., classic bridge organisms in oral biofilms. Although more closely tied to periodontal disease, *Fusobacterium* co-occurs with cariogenic taxa and can appear in early caries lesions. Its TA modules may support survival during shifts from a carious to an inflammatory milieu, potentially explaining co-aggregation and persistence across caries, periodontal disease, and even oral cancer progression [38, 48, 49].

Although *S. mutans* TA genes were not annotated through TADB3 in our dataset, established TA biology in this species is informative. Its genome encodes multiple TA modules, including a type I TA system (*fst/srSm*) consisting of a small toxic peptide and an antisense RNA antitoxin. It is the first TA module identified in streptococci [17]. The chromosomal *mazEF* and *relBE* TA loci in *S. mutans* were shown to be functional type II systems implicated in stress tolerance and antibiotic persistence [16, 18]. Intriguingly, *S. pneumoniae* (a transient oral colonizer and relative of *S. pseudopneumoniae* in our samples) also carries a relBE TA operon that enhances oxidative-stress resistance and biofilm formation [50]. This conservation of RelBE-family systems across oral streptococci suggests a common strategy for coping with the reactive oxygen and low-pH stress characteristics in dental plaque. *S. mutans* further harbors a unique three-gene TA module (SmuATR) that is induced by its competence-stimulating peptide (CSP) quorum signal. Activation of this tripartite TA operon by CSP drives a fraction of the population into persistence, demonstrating crosstalk between quorum sensing and TA-mediated dormancy [19]. Beyond isolated reports, systematic characterization of TA systems in oral bacteria remains limited. Our findings help close this gap by identifying multiple TA modules across diverse oral taxa, suggesting that both commensals and pathogens employ TA-mediated dormancy as part of their adaptive repertoire.

Functionally, most TA loci we identified belong to the RelBE/ParDE superfamily of toxins and antitoxins. These typically act as mRNA interferases (such as RelE, YoeB, and YafQ toxins) or gyrase inhibitors (ParE toxins) that can reversibly stall cell growth [8]. RelE/ParE family members are broadly implicated in replication control, survival, persistence, and biofilm formation [13]. ParDE systems, extensively studied in Enterobacteriaceae, can stabilize antibiotic-resistant plasmids and confer marked tolerance to antibiotics and heat [51]. The presence of ParE/RelE homologs in oral bacteria suggests analogous advantages, enduring transient antibiotic exposure (topical antimicrobials or prescribed systemics) and withstanding rapid temperature or pH shifts (ingested hot/cold foods, acid pulses from sugar metabolism). Consistent with this, disrupting relBE modules in *Pseudomonas aeruginosa* diminishes biofilm formation and persistence, highlighting TA loci as potential anti-biofilm targets [52]. By extension, though still speculative for the oral cavity, selectively inhibiting certain TA systems in dental plaque could weaken biofilm resilience and enhance caries interventions.

Several specific TA components merit particular attention. We detected transcripts for the toxin YafQ (of the DinJ-YafQ Type II TA pair) without its cognate antitoxin DinJ. DinJ/YafQ modules are common in Gram-positive bacteria like *Lactobacillus*, where YafQ is a ribosome-dependent endoribonuclease that DinJ usually neutralizes. YafQ can become unstable or degraded in the absence of DinJ, yet under stress, it can persist long enough to inhibit translation and transiently arrest growth [53, 54]. Detecting YafQ alone may reflect partial acquisition or truncation of the locus (e.g., via horizontal gene transfer) or differential regulation with toxin transcript levels temporarily outpacing antitoxin expression. This imbalance warrants further investigation, as it could indicate alternative buffering mechanisms or a role for YafQ in interspecies interactions. We also identified members of the VapBC family (toxin VapC with antitoxin VapB). VapC toxins are RNases that typically target tRNA or rRNA. VapBC modules are prevalent in pathogens like *Haemophilus influenzae*, where they enhance survival during infection [55]. Their presence in oral taxa suggests analogous functions in enduring fluctuating pH, oxidative stress, and nutrient pulses within dental biofilms.

Environmental cues are central to TA activation. Nutrient limitation, DNA damage, oxidative stress, and quorum-sensing signals can destabilize antitoxins, often via protease activity, liberating the toxin [9, 11]. In the highly dynamic oral niche, microbes encounter repeated triggers: acid pulses from sugar fermentation, oxidative bursts from host immunity, and intermittent exposure to agents such as fluoride or chlorhexidine. We propose that oral bacteria use TA systems to sense and respond to these stresses by entering a transient semi-dormant state, preserving a reservoir of survivors when metabolically active cells are compromised. A recent review supports this model, noting that TA systems respond to diverse environmental signals to modulate competence, biofilm formation, and persistence [56]. In dental caries, TA-driven persister formation during acid shocks or post-treatment intervals could facilitate rapid biofilm rebound, helping explain recurrent lesions and the difficulty of fully eradicating pathogenic consortia when a protected subpopulation is maintained through TA activation.

We acknowledge some limitations of our analysis. Our TA gene annotations relied on TADB 3.0, a comprehensive database of experimentally validated and predicted TA loci [28]. While TADB covers tens of thousands of genomes, it may not include every oral bacterium (many oral taxa remain less studied), so some TA genes could have escaped detection. In particular, novel TA systems or those in species without reference genomes might be missed. As a secondary analysis of existing datasets [20, 21], our conclusions are constrained by the design of those studies and sequencing depth. Low-abundance TA transcripts or context-specific activation may have gone undetected. We also inferred function from expression and did not measure toxin or antitoxin protein activity directly, so some TA loci may have been transcribed without physiological effect. Even so, the concordance with established TA biology and prior reports supports the credibility of our interpretations.

Our study opens several avenues for future research. Controlled in vitro experiments could validate the function of specific oral TA systems, for example, knocking out or overexpressing a *Leptotrichia* or *Streptococcus* TA gene to test effects on biofilm formation, acid tolerance, or antibiotic persistence. Monitoring TA gene expression in real time during pH stress or sugar exposure could directly link environmental triggers to TA activation in oral bacteria [57]. Exploring TA systems beyond Type II in the oral microbiome would also be valuable. Types I and III-VIII, which involve various RNA antitoxins or other mechanisms [15, 35], remain almost completely uncharted in dental plaque. Unraveling those could reveal additional layers of microbial interaction and competition. Another intriguing direction is to examine TA systems as potential therapeutic targets in dentistry. If certain TA modules are essential for the survival of cariogenic bacteria under stress, small molecules or enzymes that disrupt the toxin-antitoxin balance might render the biofilm more susceptible to conventional treatments. This concept, sometimes called “anti-persister” or “anti-biofilm” therapy, has gained traction in medical microbiology [58], and the oral cavity could similarly benefit from such innovative approaches.

## Conclusion

Our refined analysis reveals that oral microbes broadly express TA systems and modulate them across healthy, caries, and treatment states. These stress-adaptive regulators, previously underappreciated in the oral niche, likely influence community stability and therapeutic resistance. Their contributions to plaque resilience and disease recurrence warrant further investigation and may present future targets for precision strategies against biofilm persistence.

## List of Abbreviations

TA: Toxin-Antitoxin
Fl: Fluoride
Fl-Ar: Fluoride-Arginine
CA: Caries-Active
CAa: Caries-Active ANCL (Active non-cavitated caries lesions)
CAi: Caries-Active INCL (Inactive non-cavitated caries lesions)
CAs: Caries-Active Sound Dental Surfaces
CI: Caries-Inactive
CF: Caries-Free
ANCOM-BC: Analysis of Compositions of Microbiomes with Bias Correction
HUMAnN: Human Microbiome Project Unified Metabolic Analysis Network
TADB: Toxin-Antitoxin Database
VFDB: Virulence Factor Database

## Declarations

### Ethics approval and consent to participate

Not applicable.

### Consent for publication

Not applicable

### Availability of data and materials

The datasets analyzed during the current study were from the BioProject (https://www.ncbi.nlm.nih.gov/bioproject/) repository – PRJNA712952 (www.ncbi.nlm.nih.gov/bioproject/PRJNA712952) [20] and PRJNA930965 (https://www.ncbi.nlm.nih.gov/bioproject/PRJNA930965) [21].

All the analyzed data and codes are available in the GitHub - https://github.com/biocoms/ta_systems_oral_mt

The source code is also available in https://doi.org/10.5281/zenodo.16763308

### Author’s contributions

SVR – Designed the study, performed all the downstream analysis, visualization, wrote the manuscript and GitHub documentation.

PS – Perfomed analysis of the metatranscriptomics pipeline and wrote that part of the GitHub documentation

EZ – Supervised and designed the study, validated the results, revised and reviewed the manuscript and documentation.

All authors read and approved the final manuscript.

## Acknowledgements

Not applicable

