## supplementary figures for "Metatranscriptomic analysis reveals toxin-antitoxin system shifts in caries-associated oral microbiomes"

**Figure S1. Volcano plots of differential expression analysis of condition comparisons in the Dieguez Dataset**

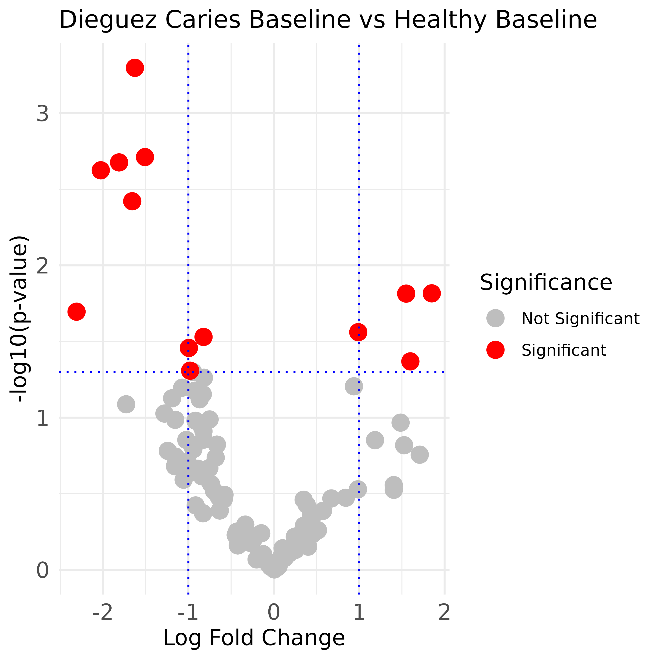

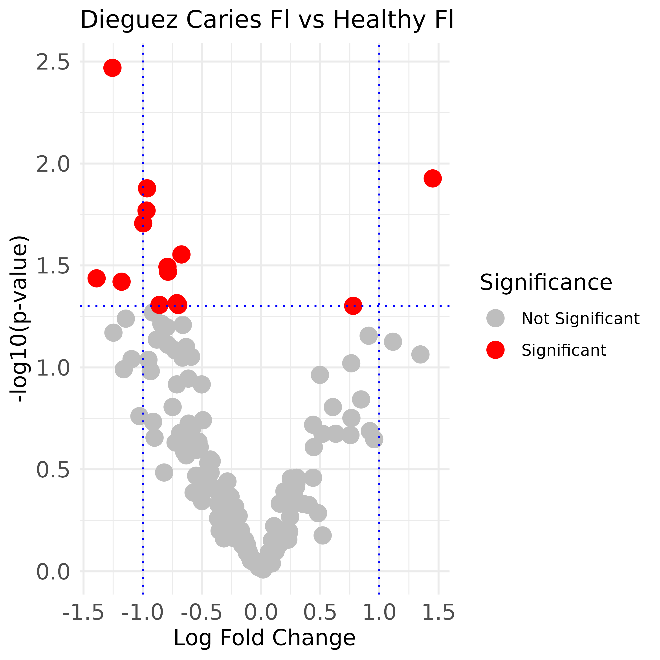

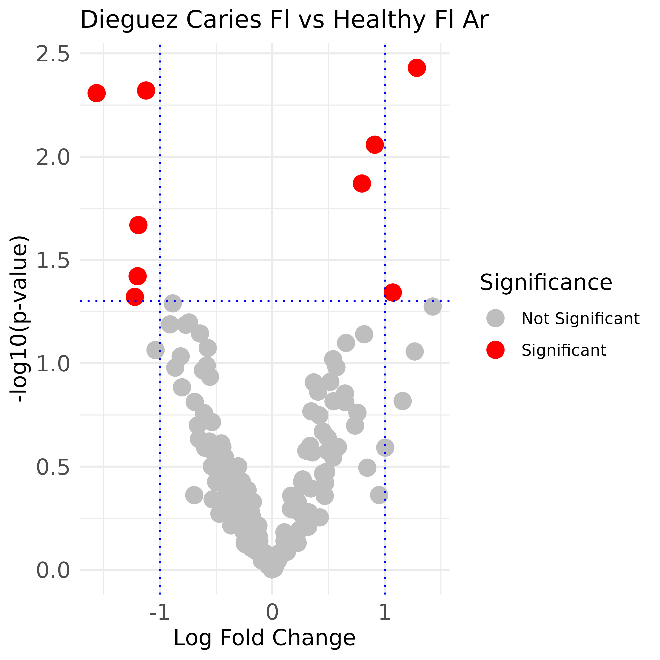

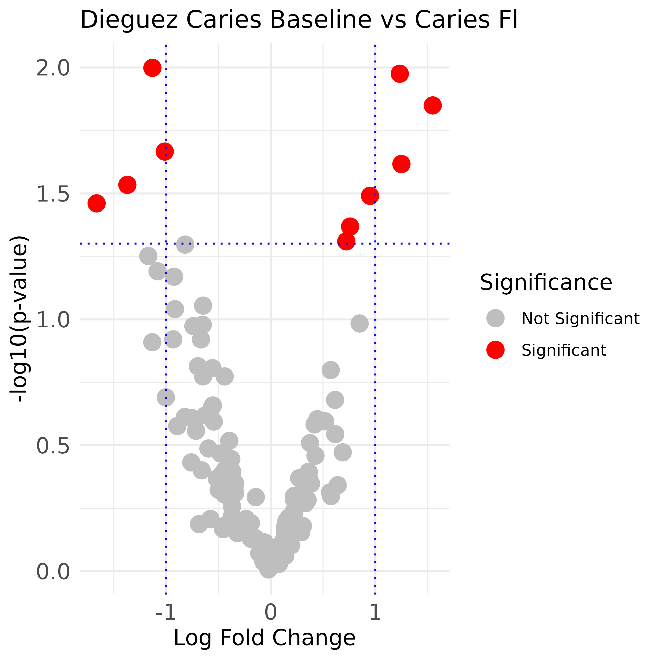

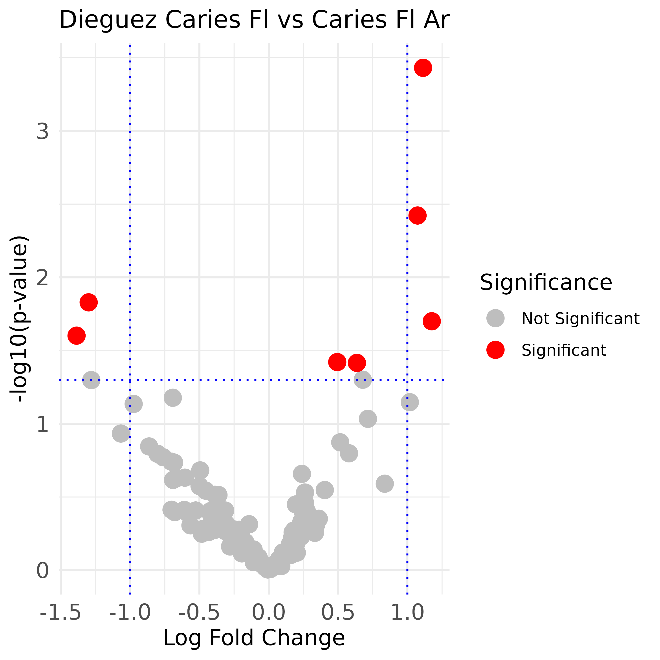

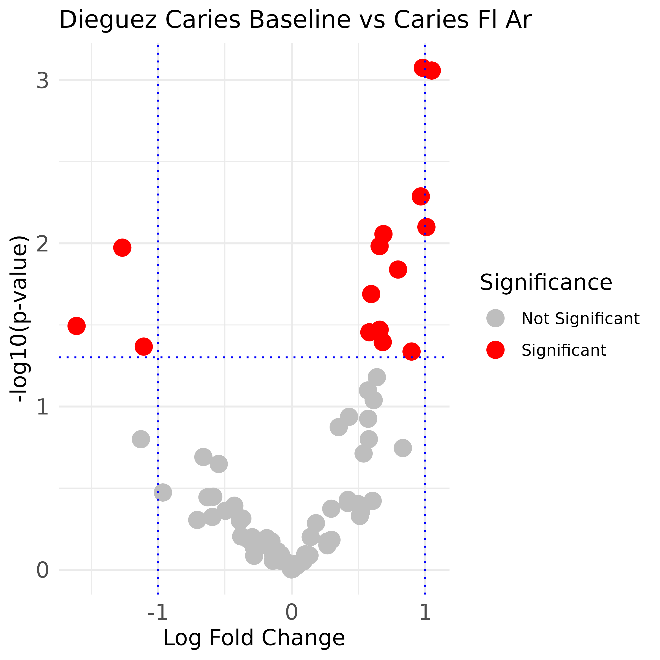

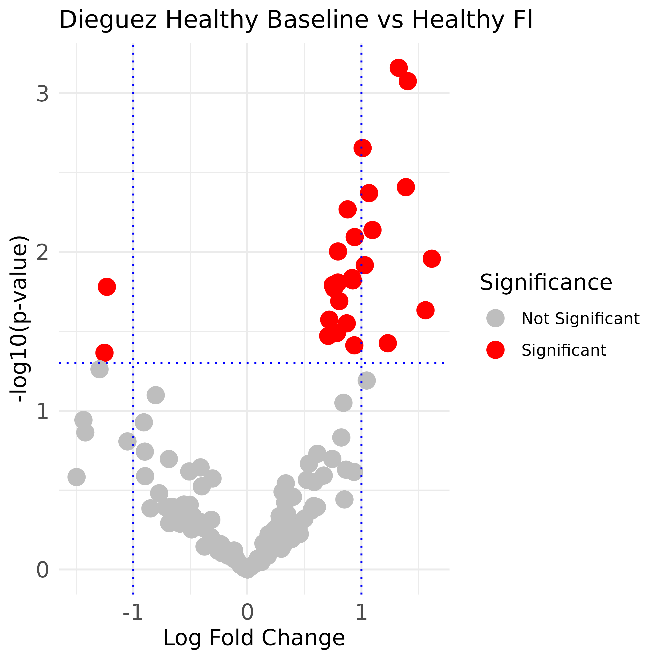

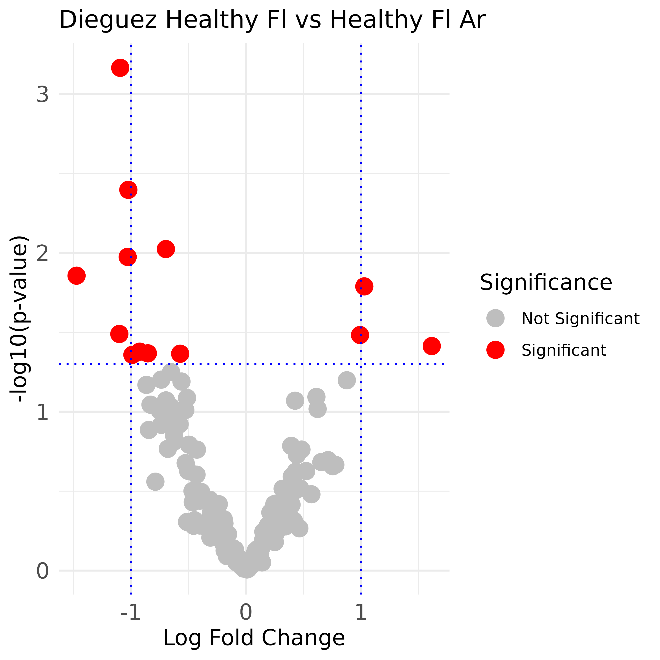

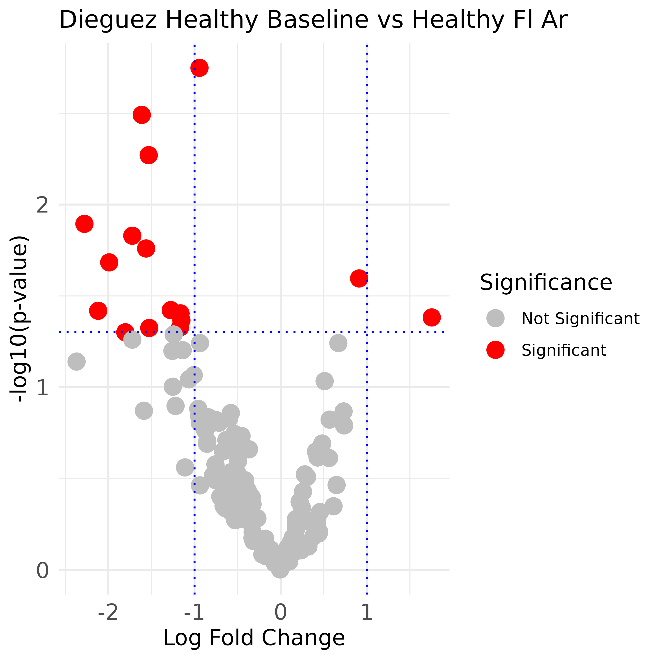

**Figure S2. Volcano plots of differential expression analysis of condition comparisons in the Ev Dataset**

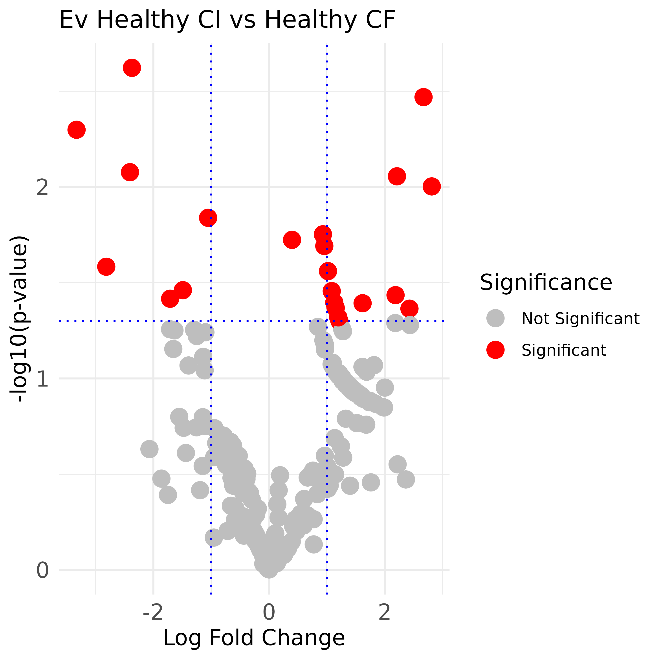

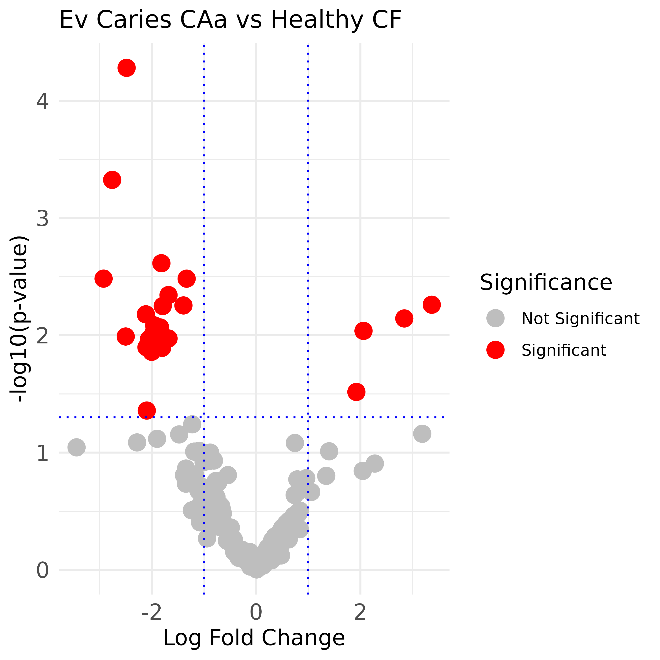

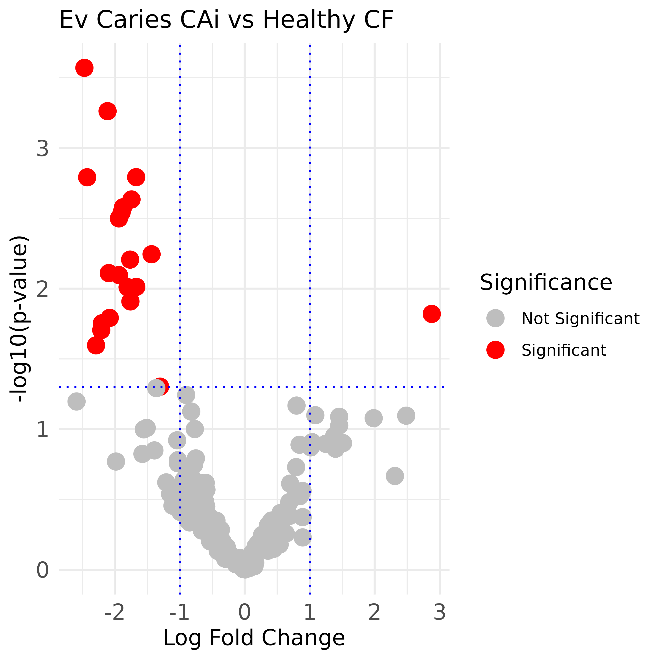

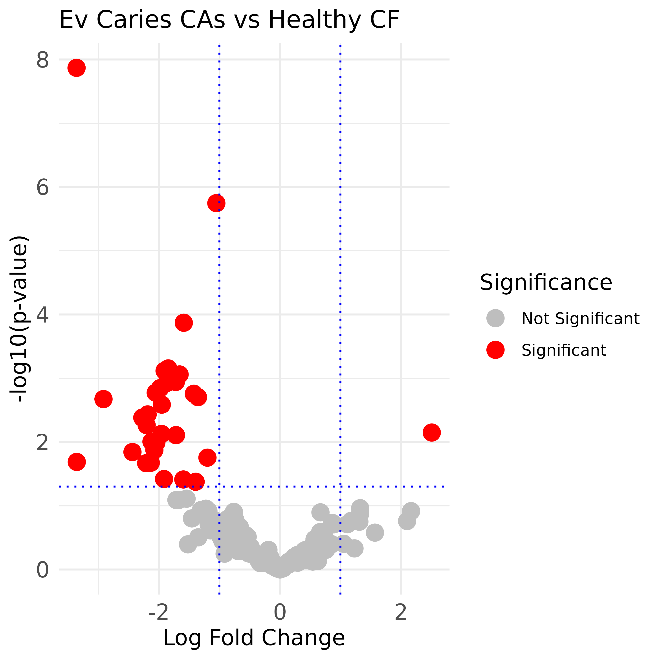

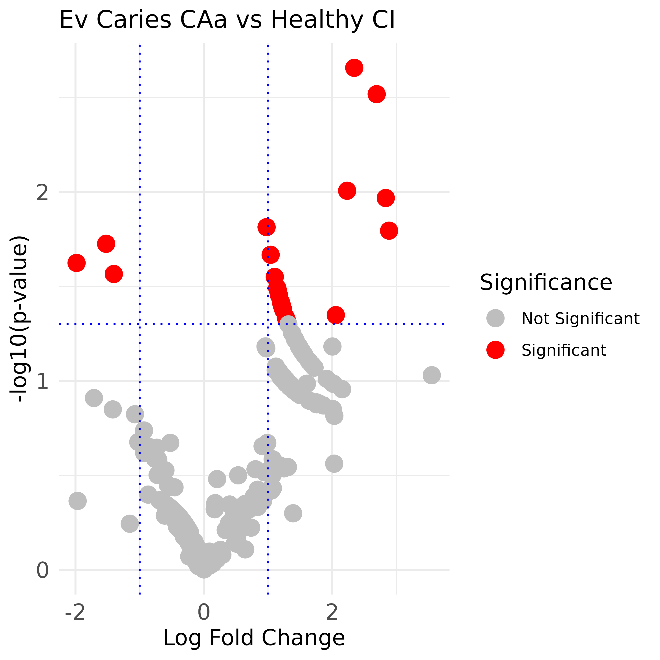

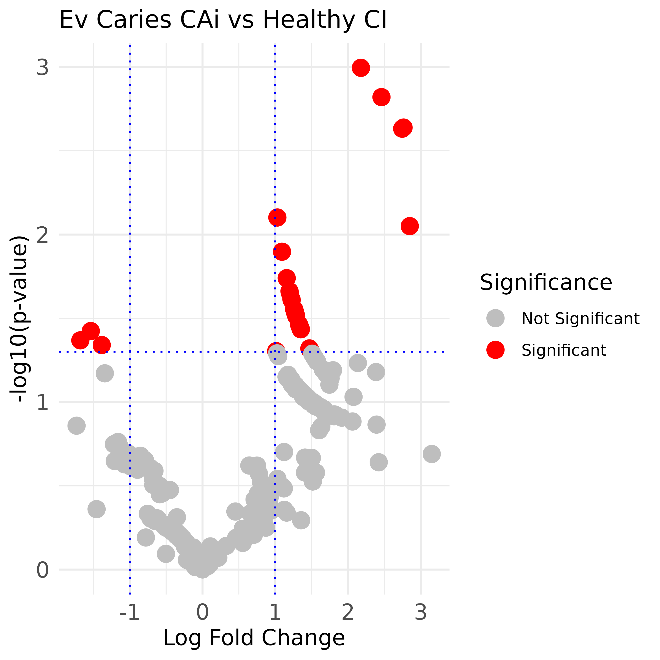

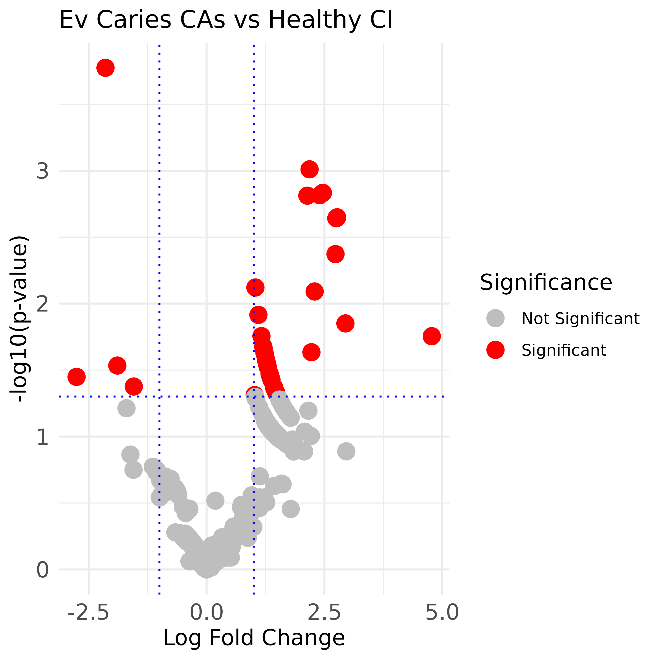

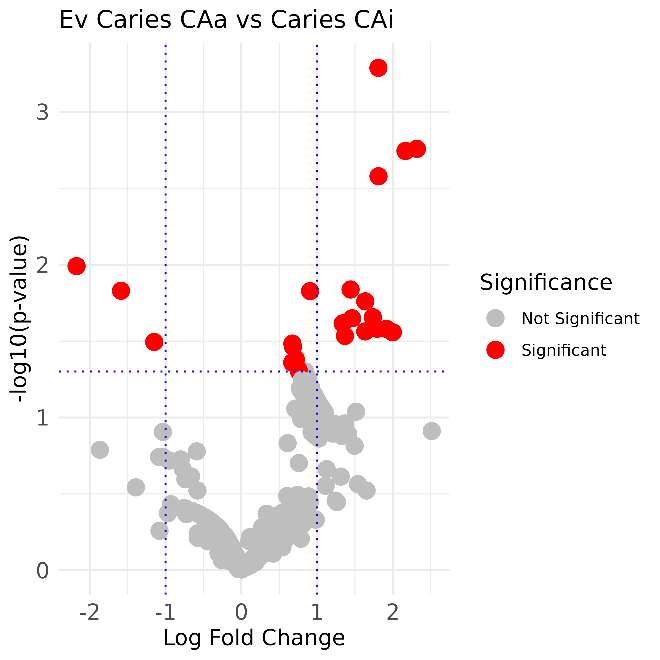

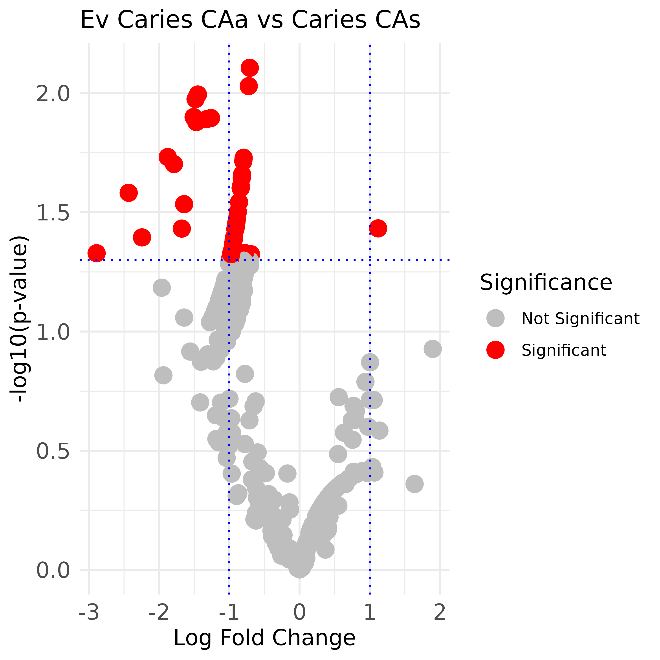

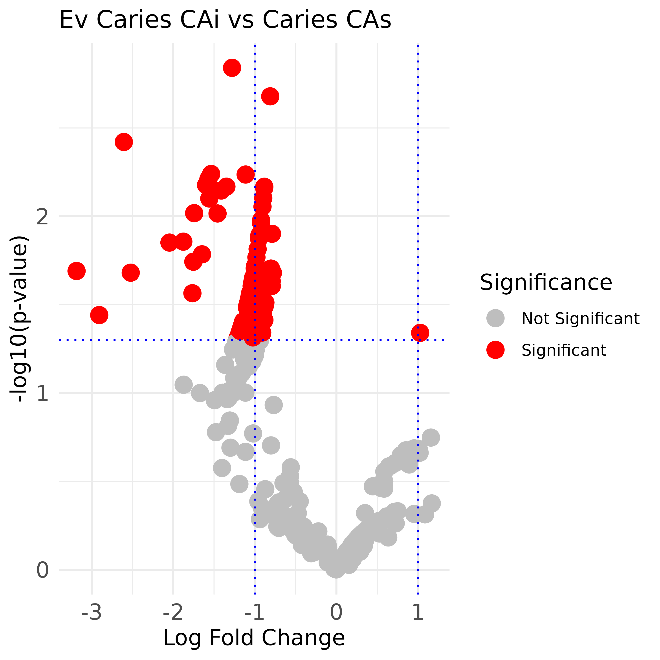

**Figure S3. Heatmaps of differential expression analysis of condition comparisons in the Dieguez Dataset**

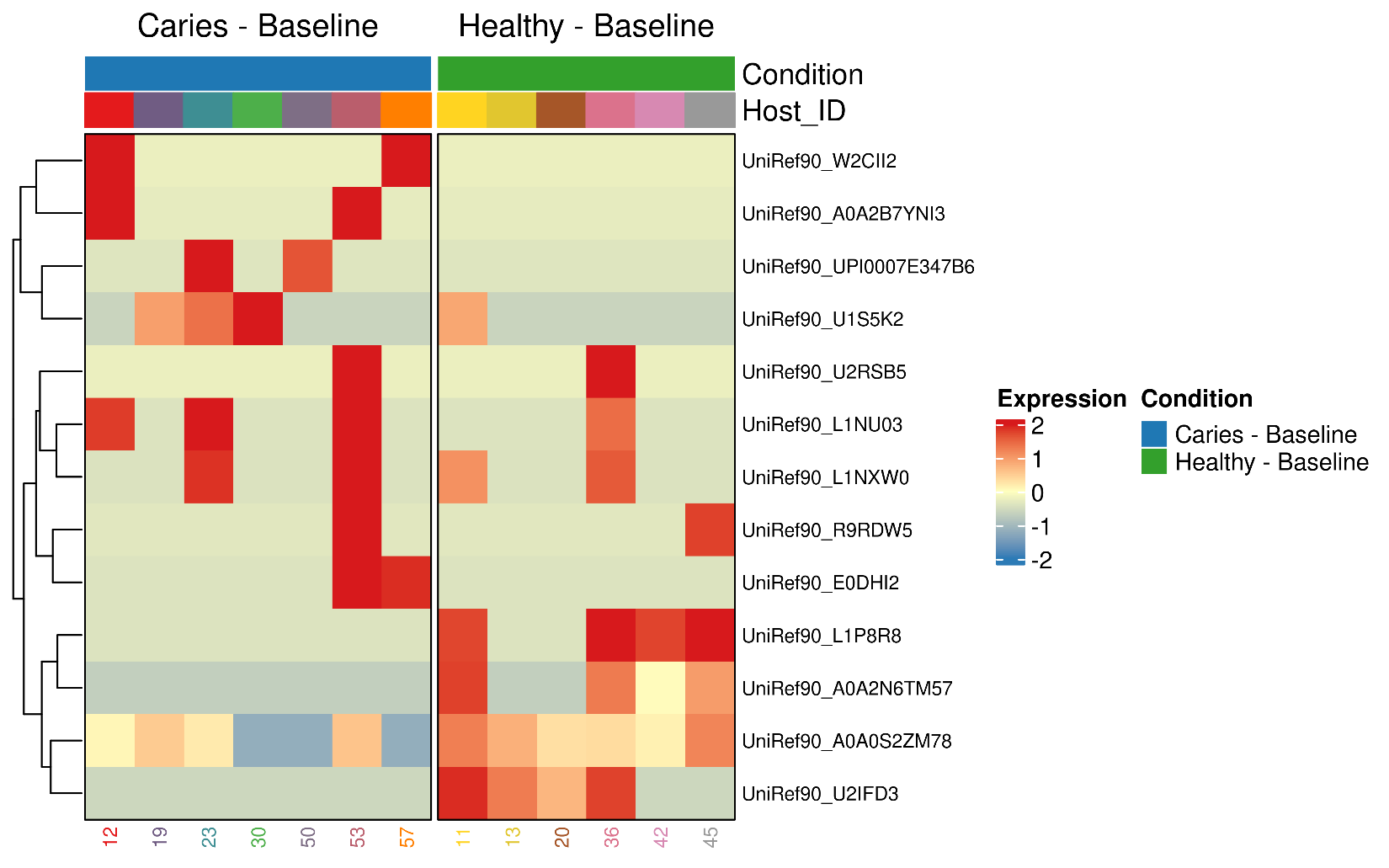

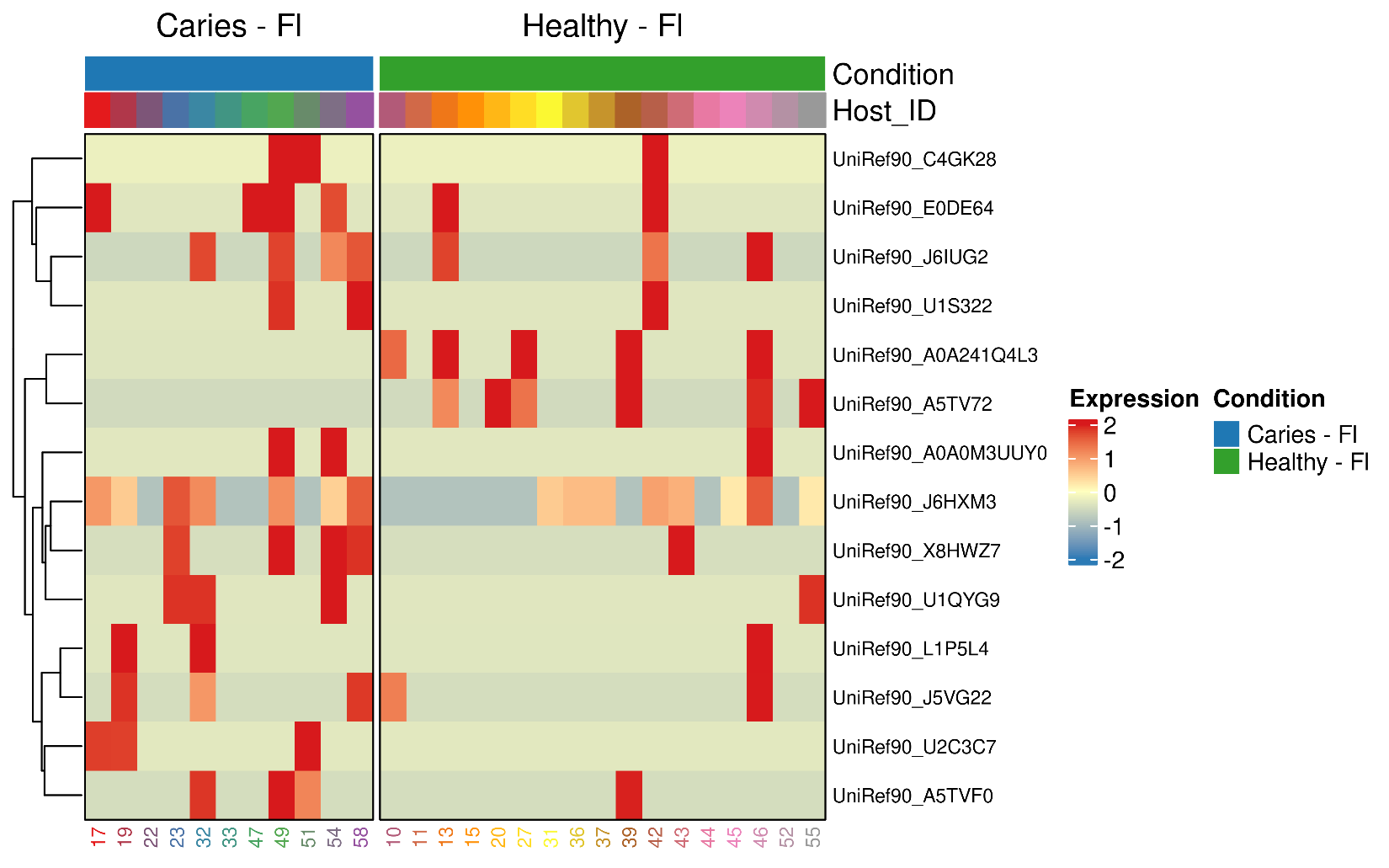

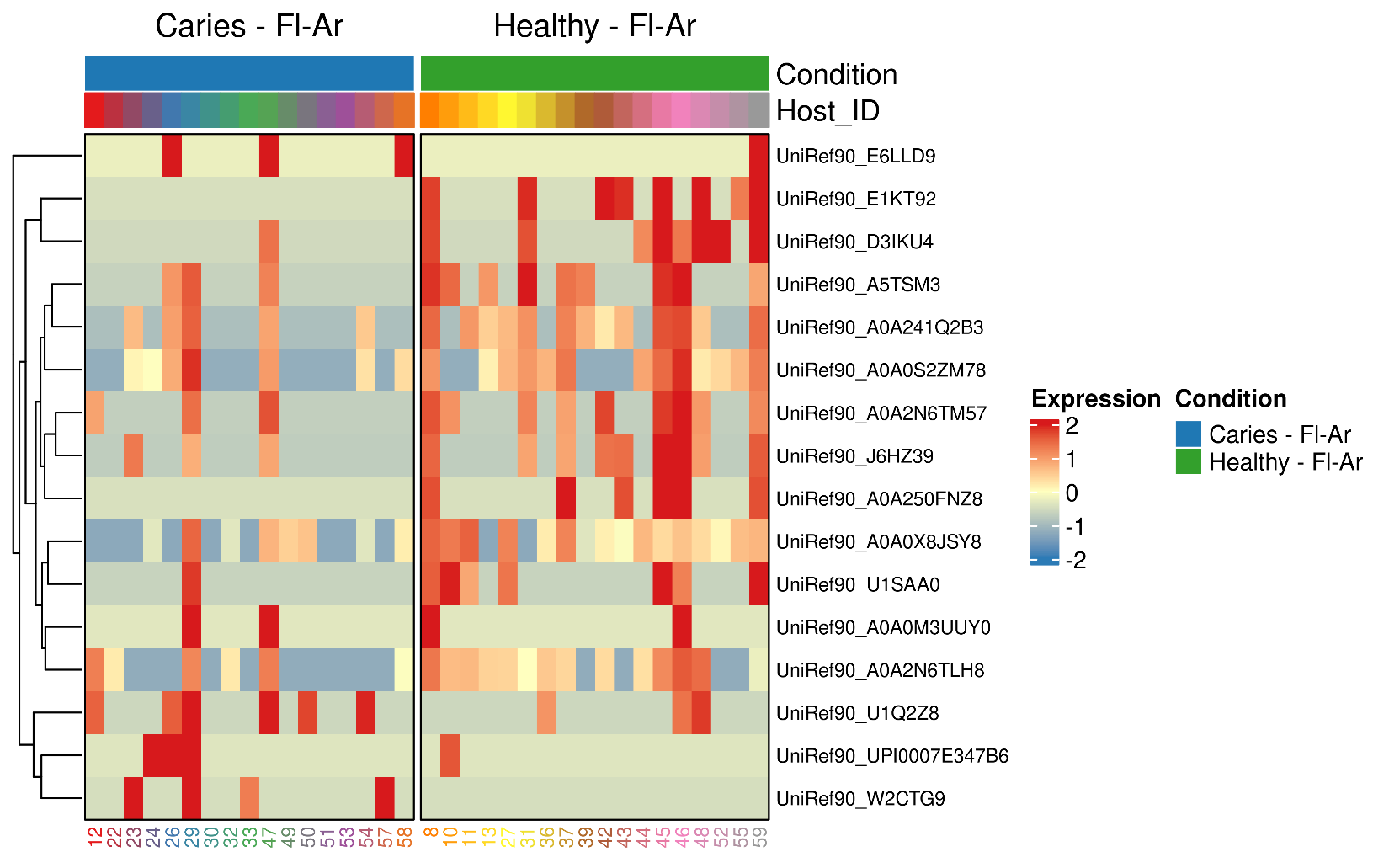

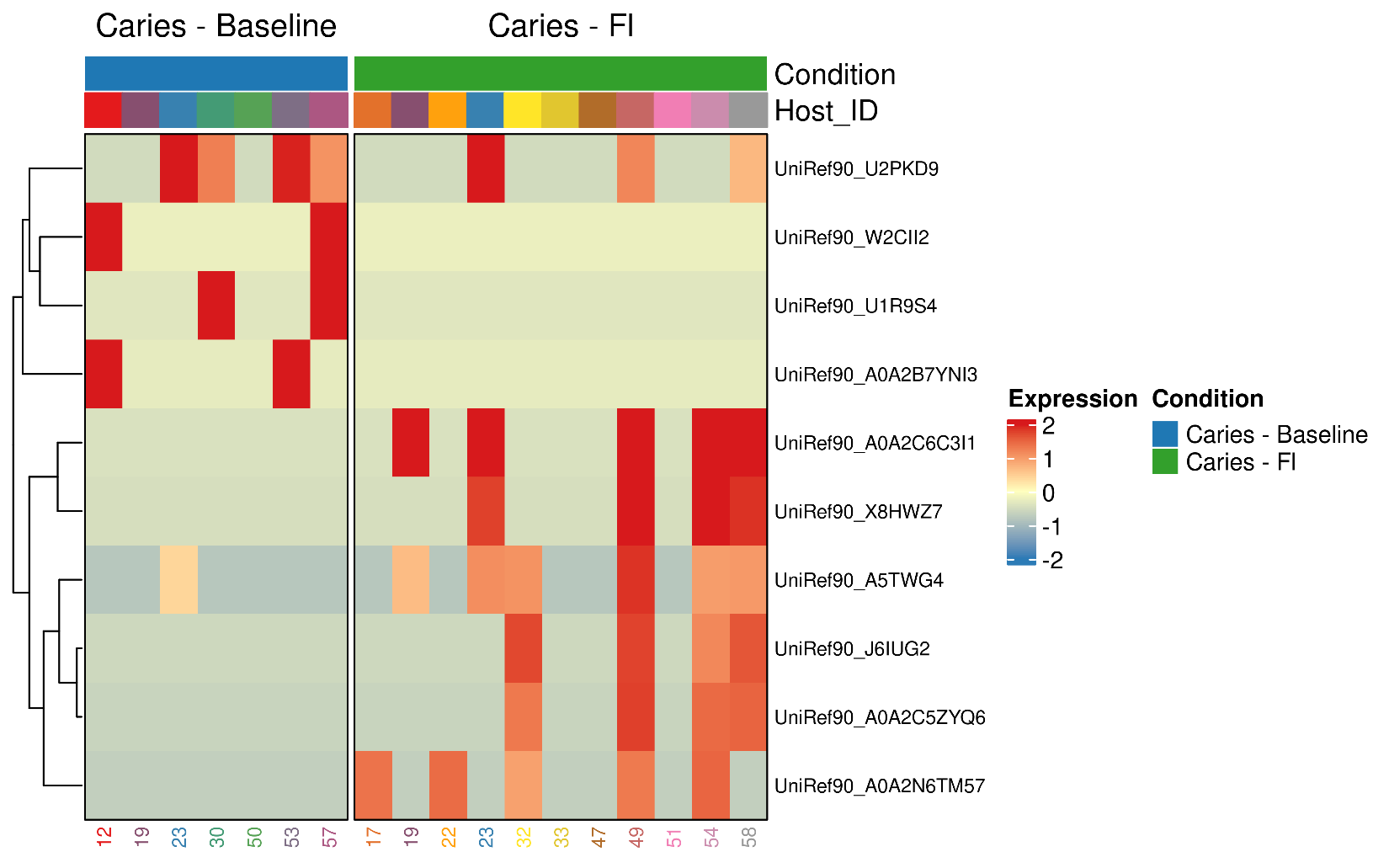

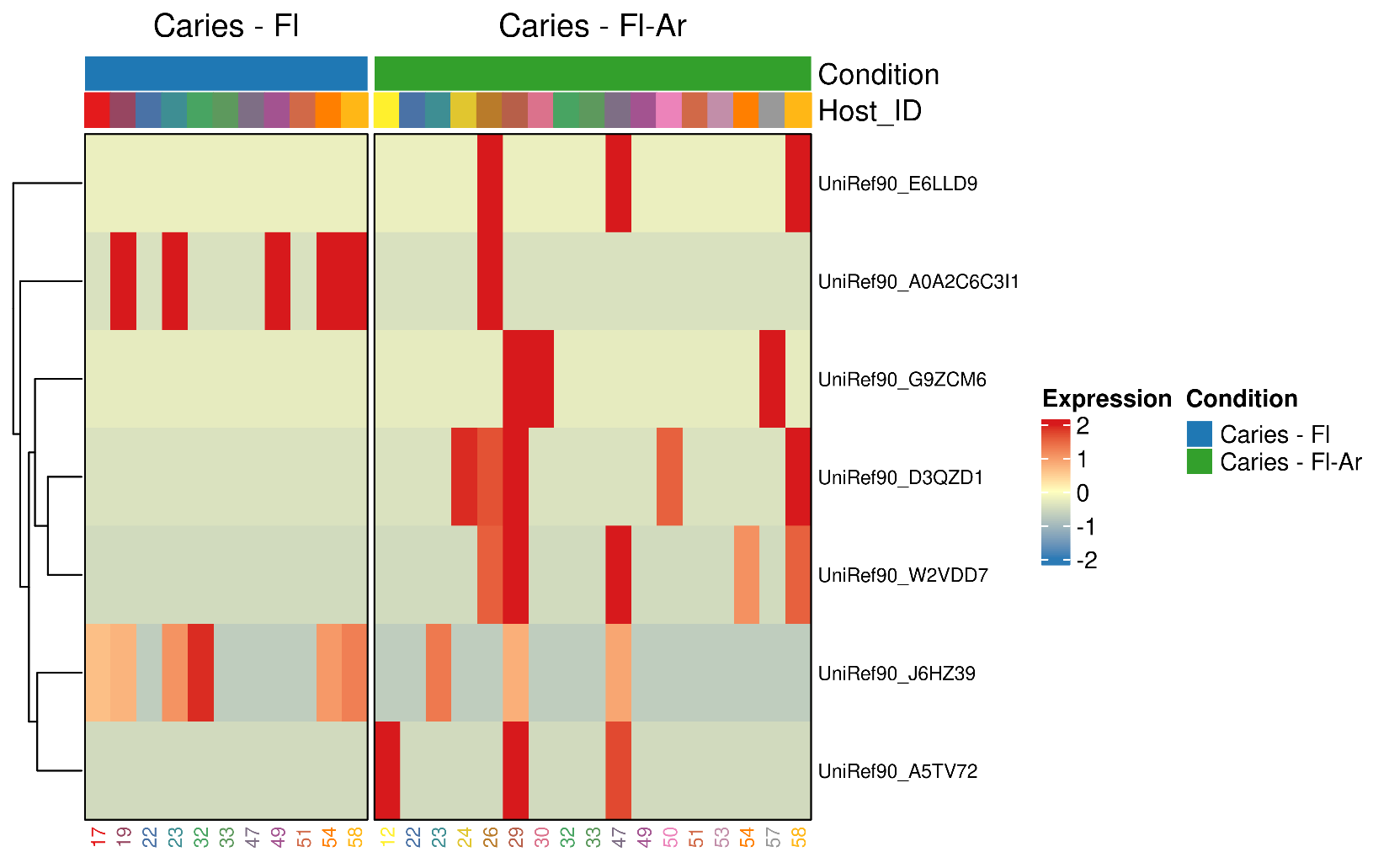

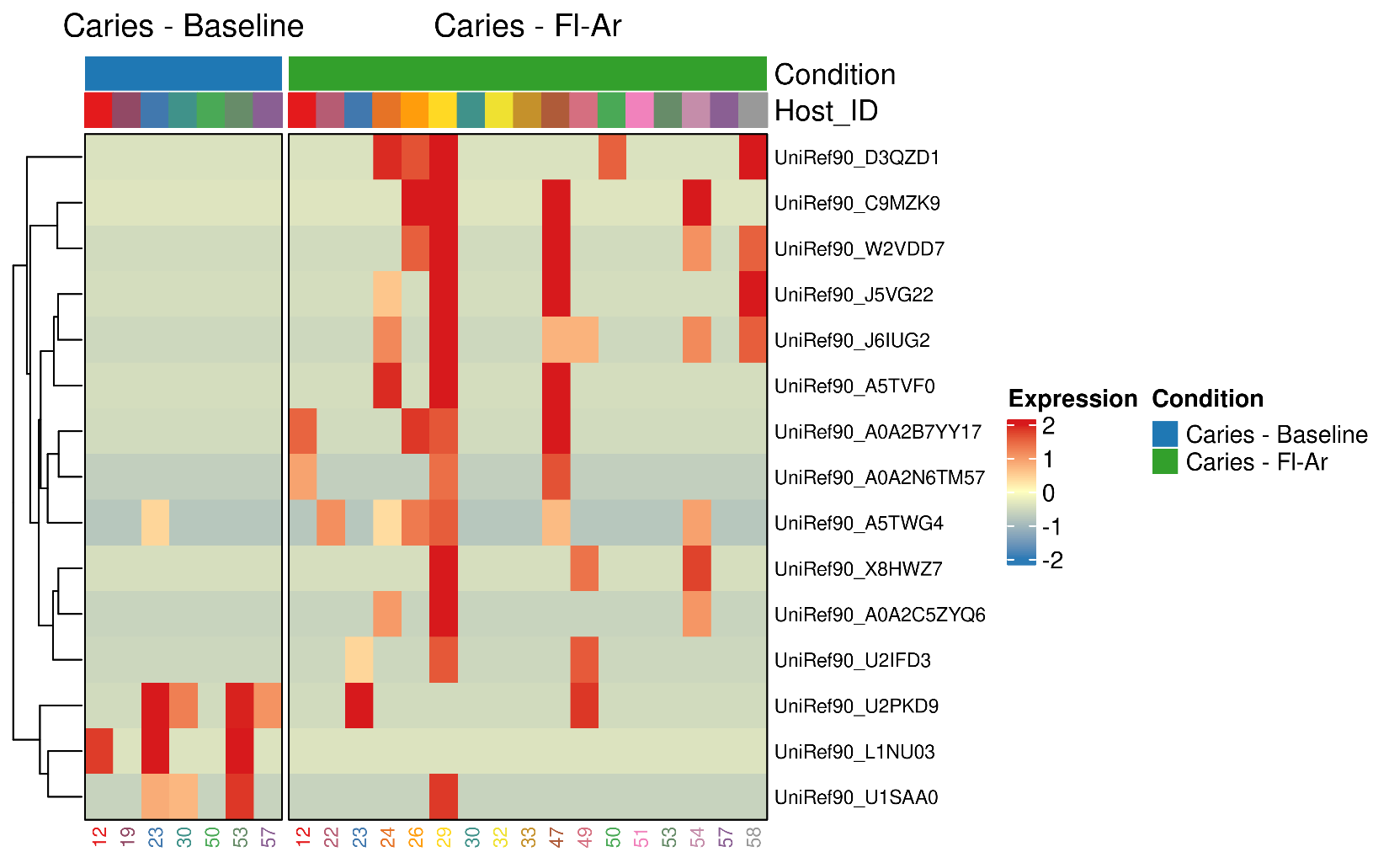

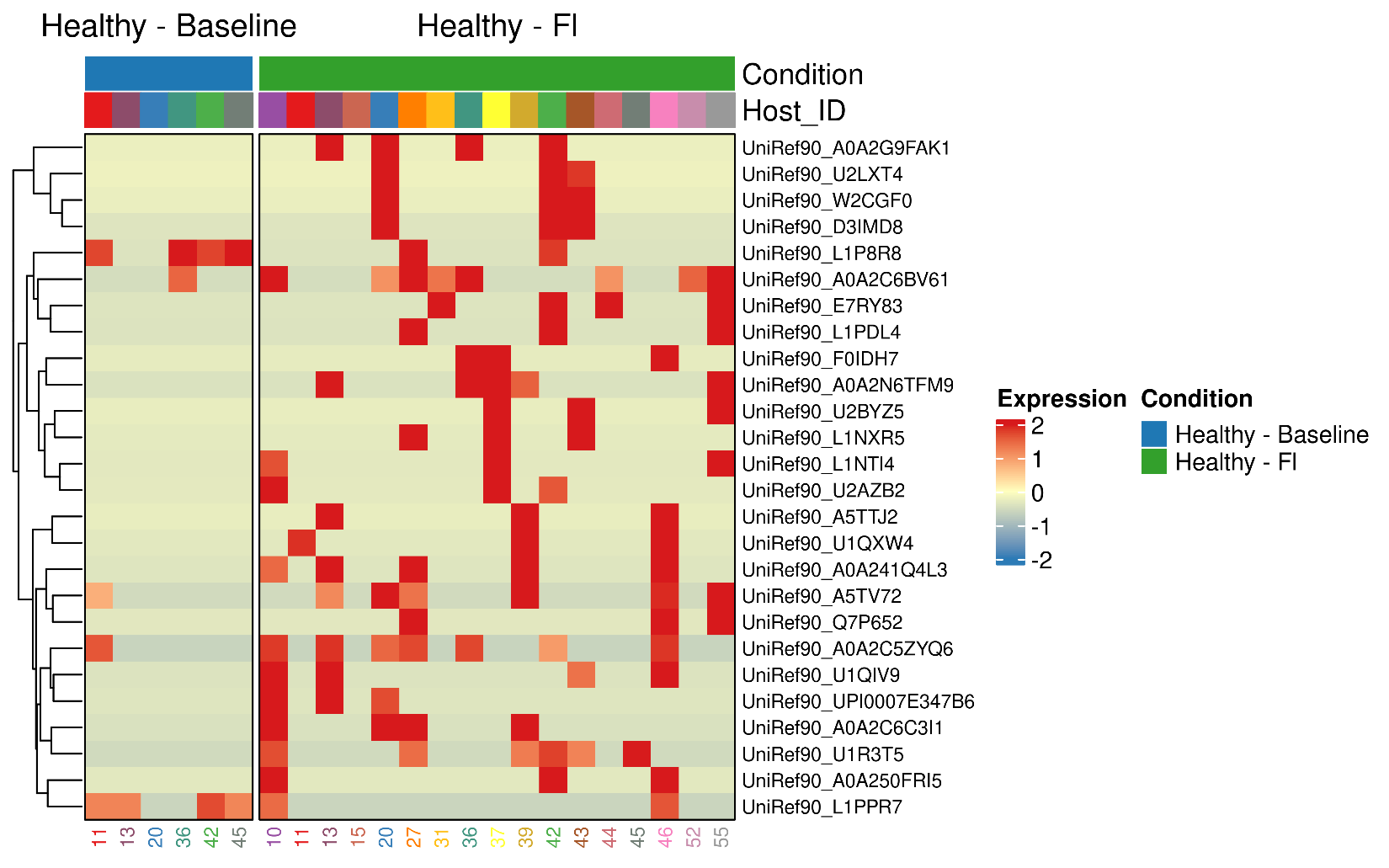

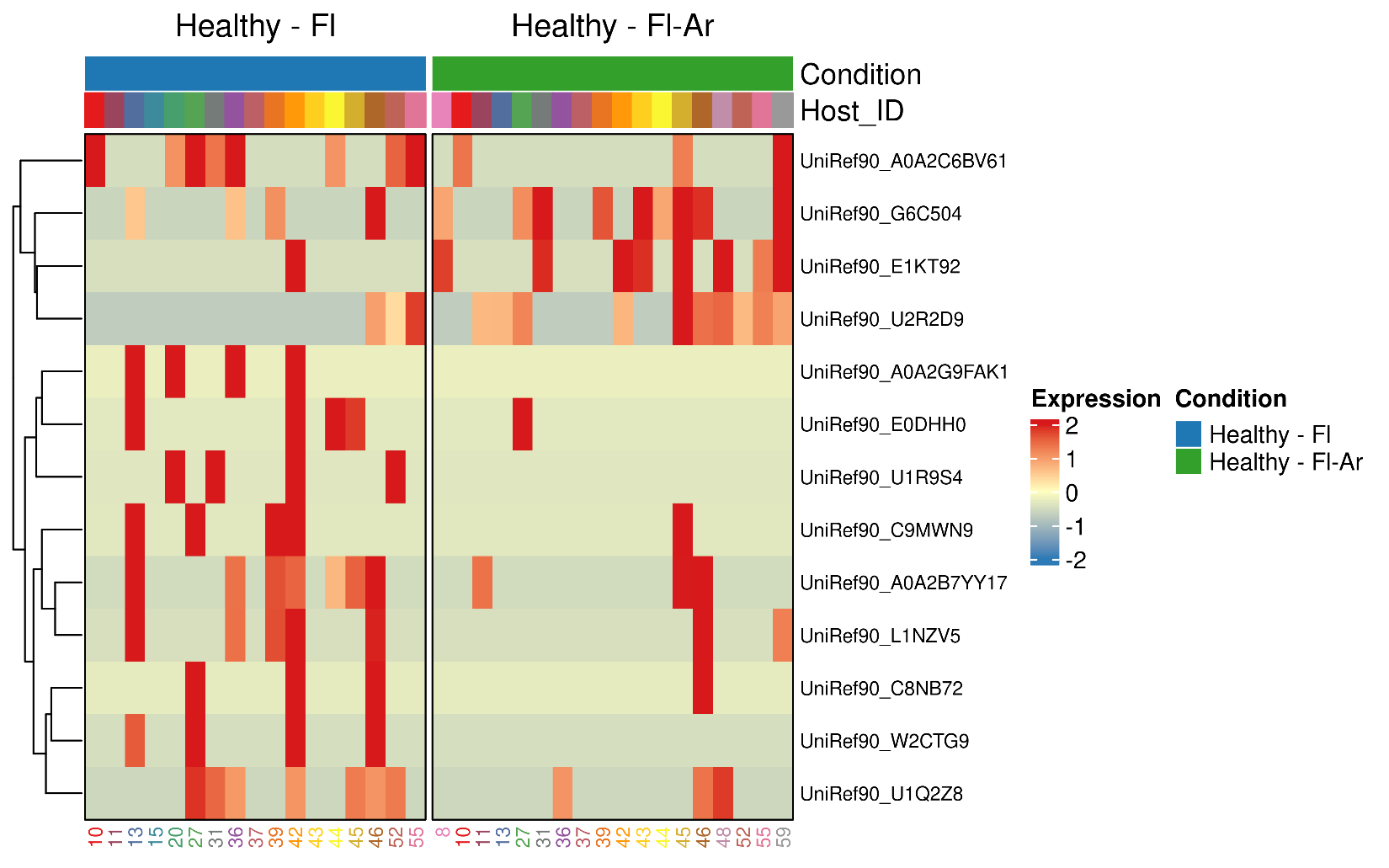

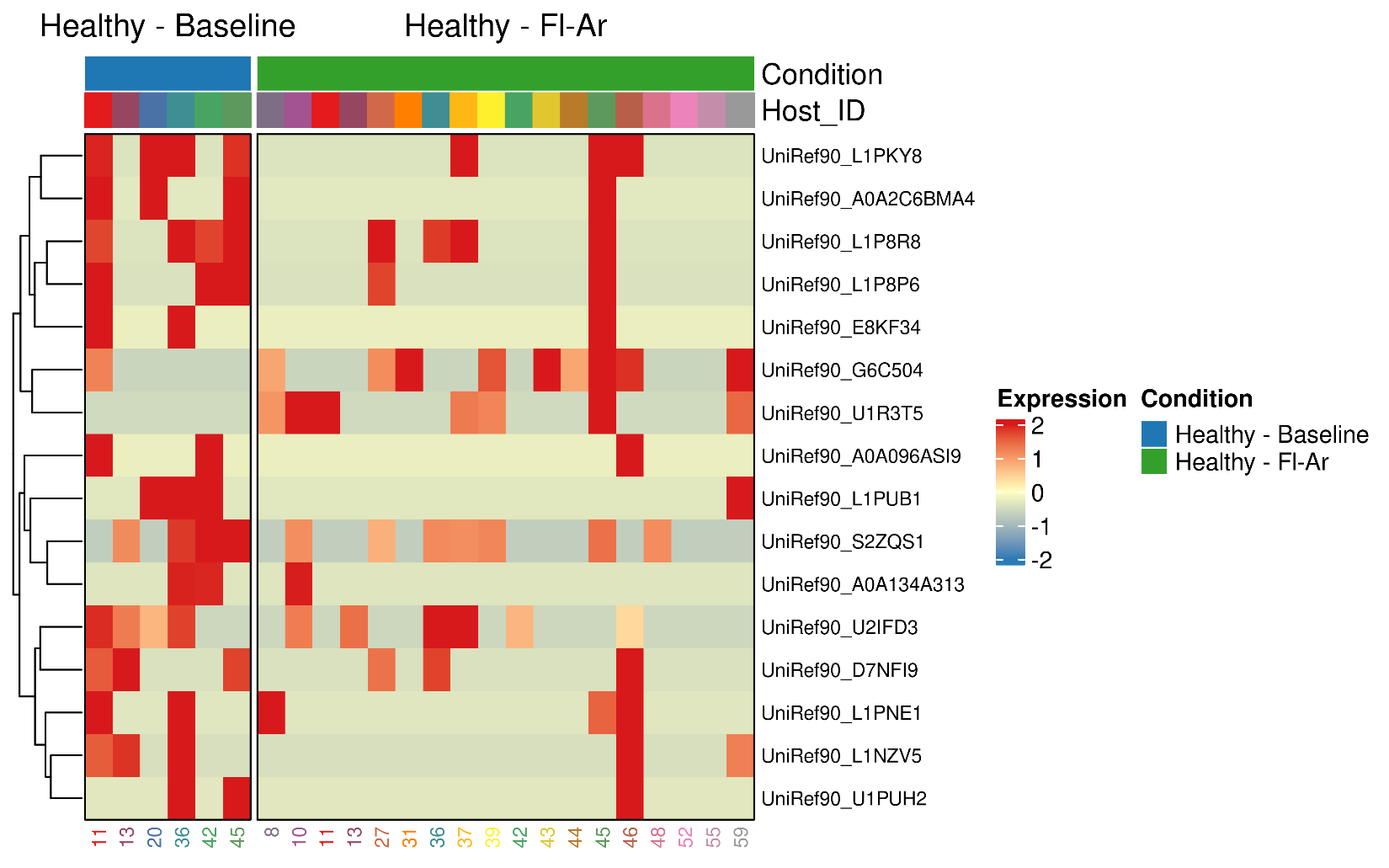

**Figure S4. Heatmaps of differential expression analysis of condition comparisons in the Ev Dataset**

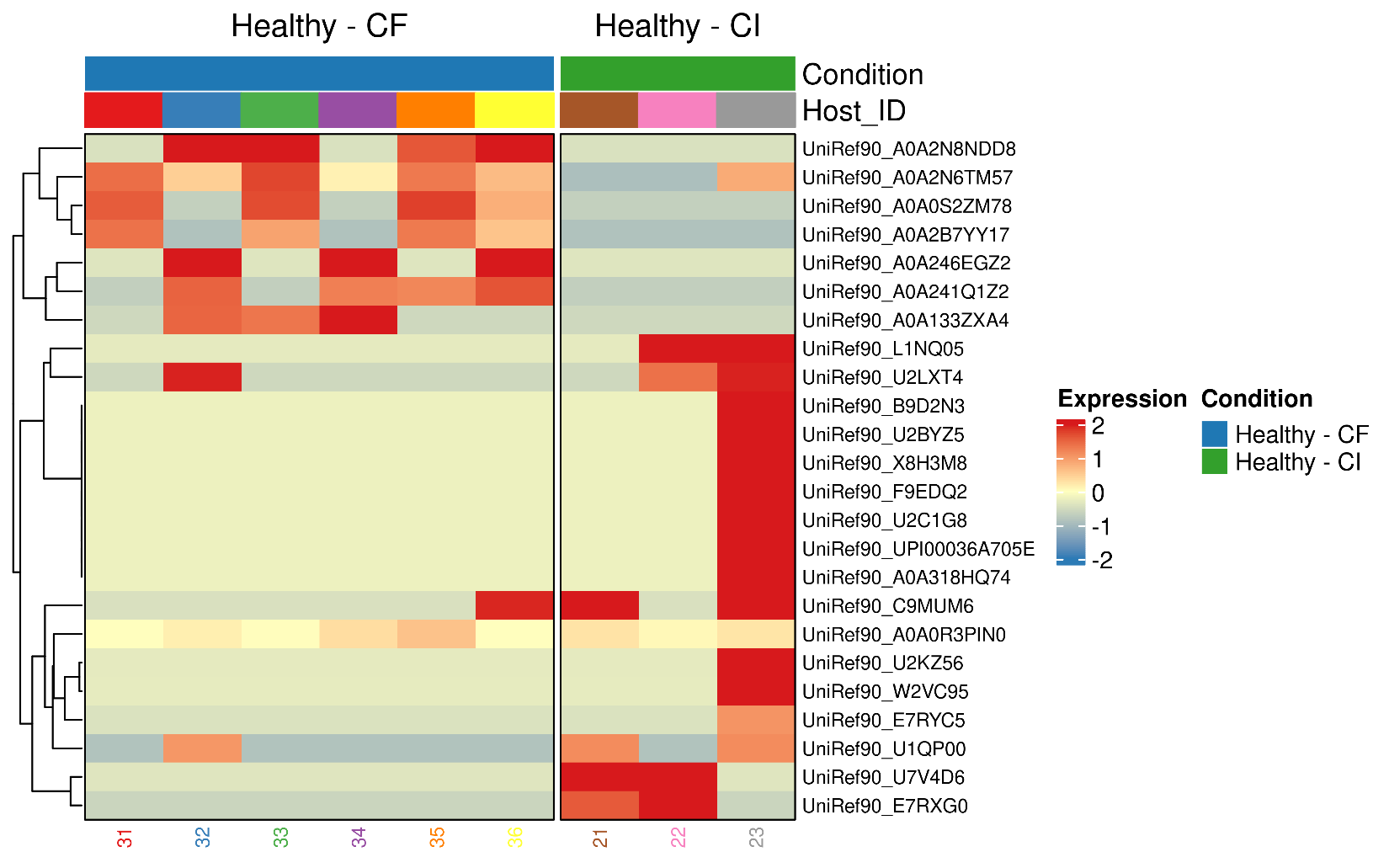

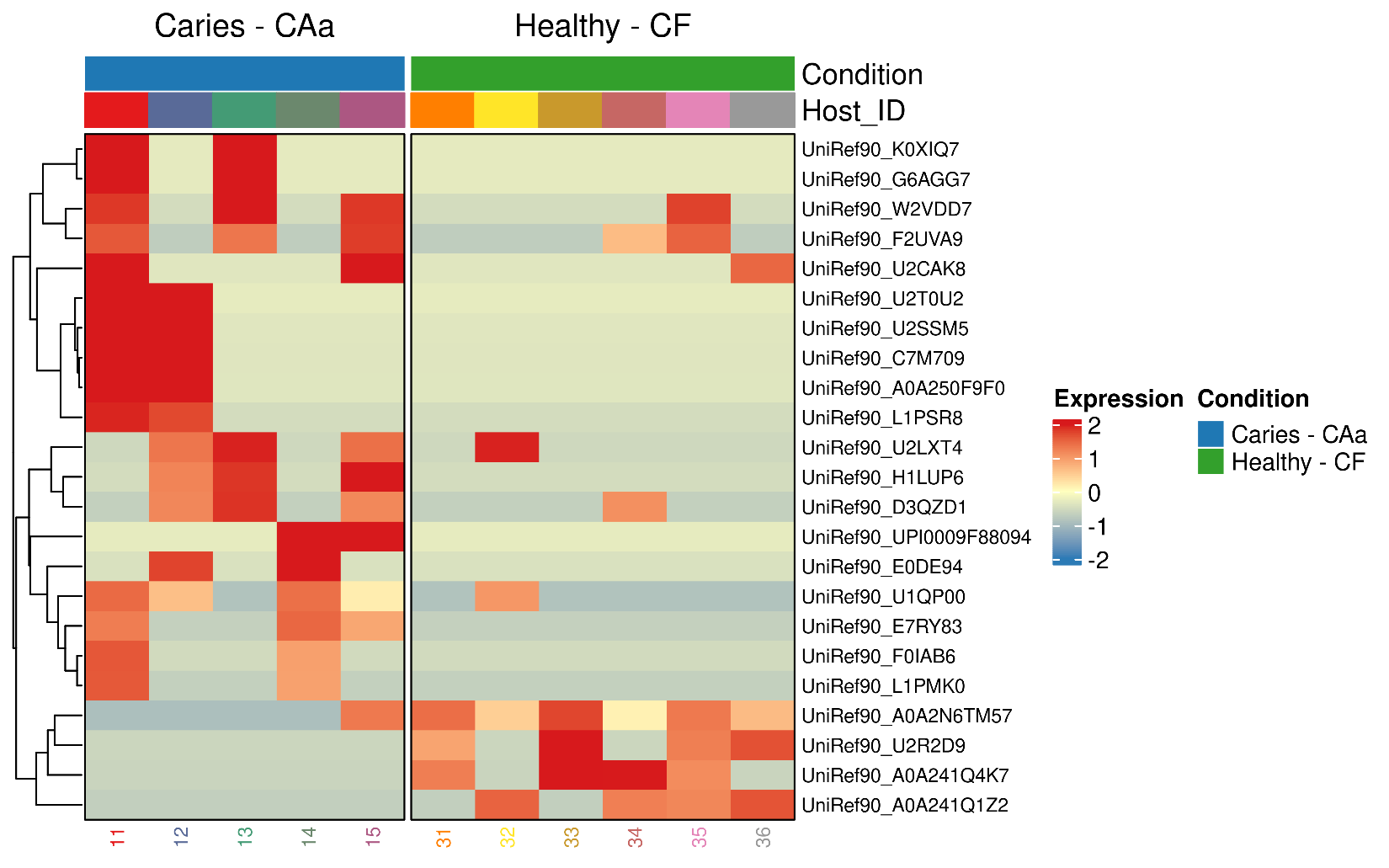

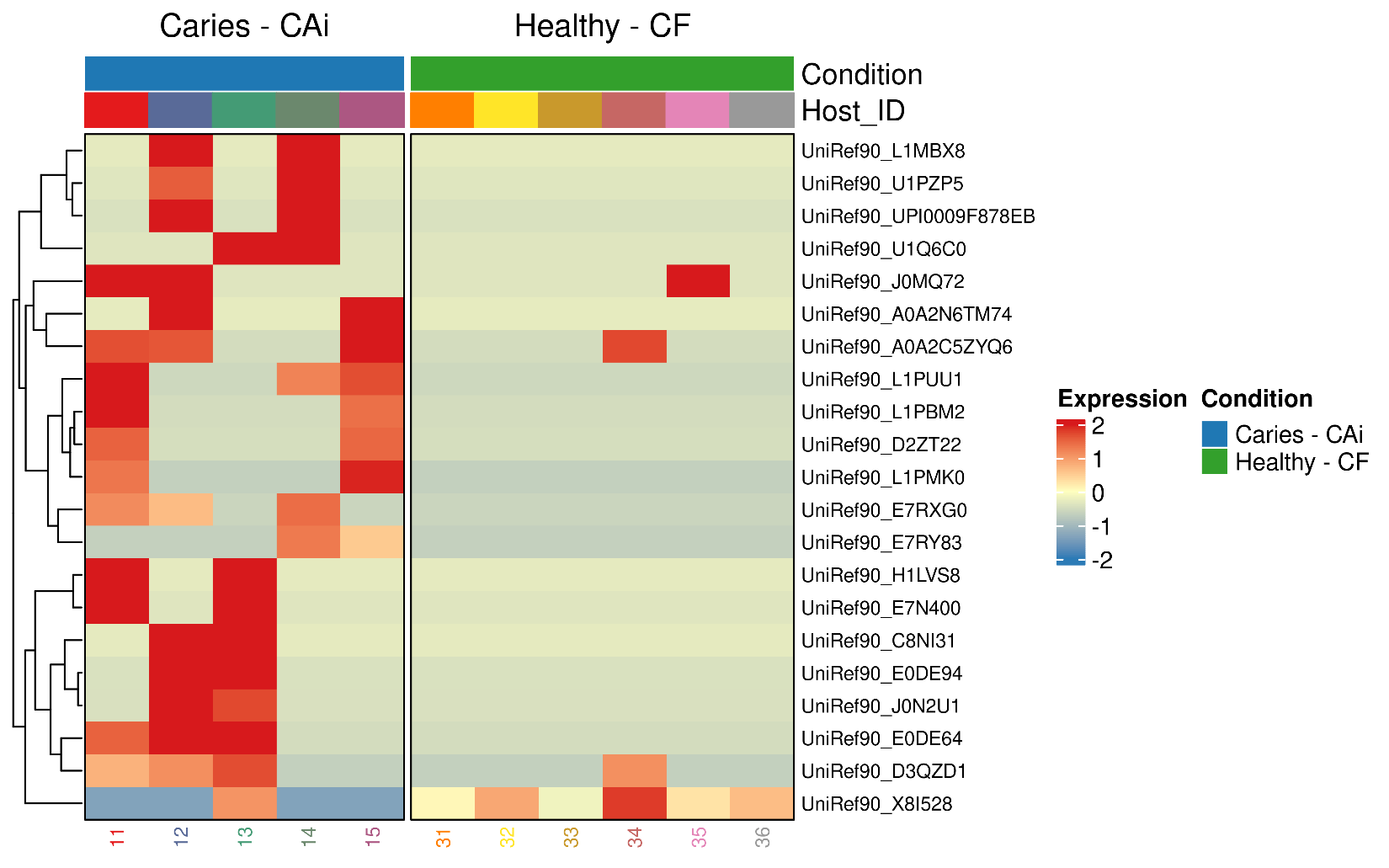

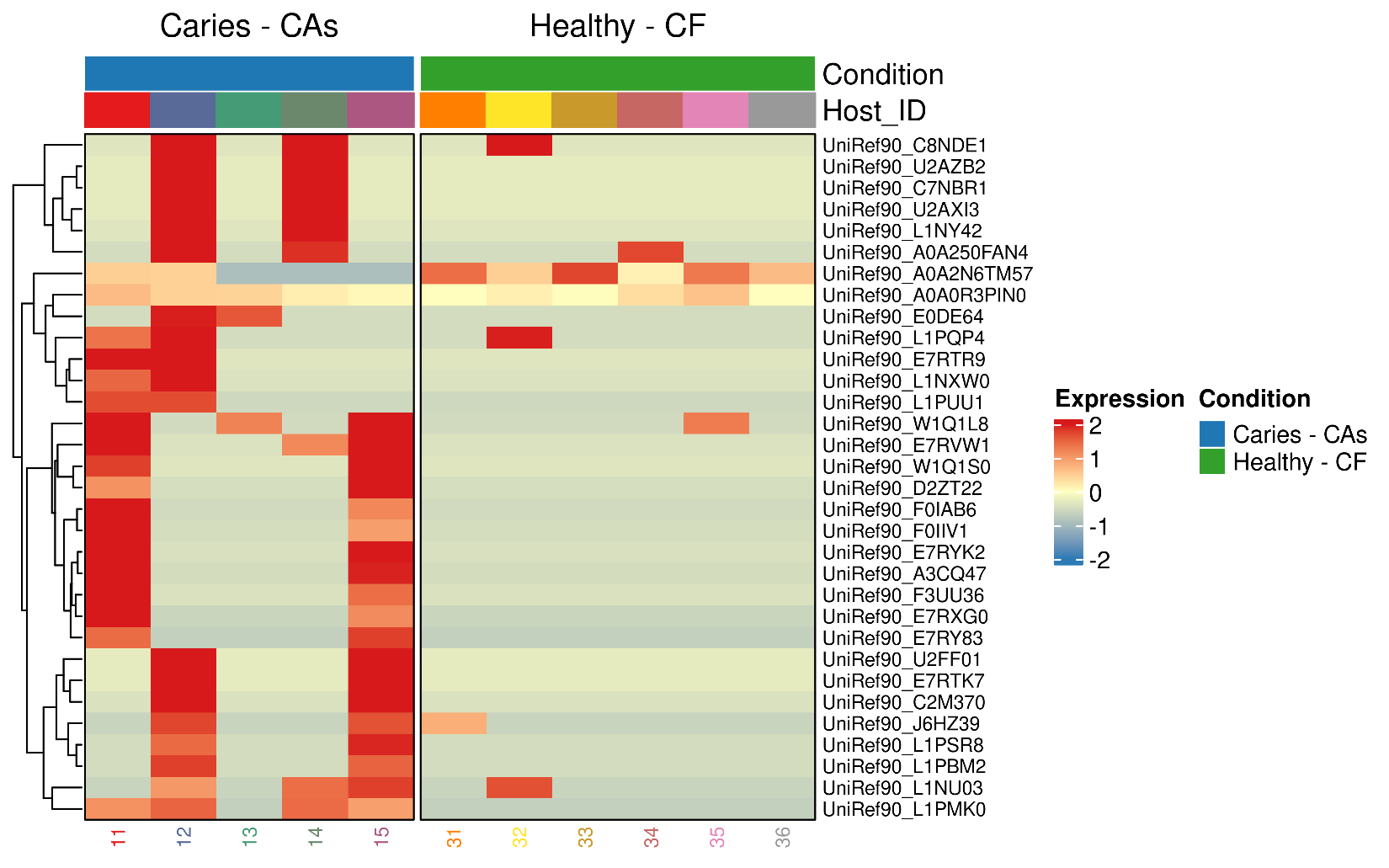

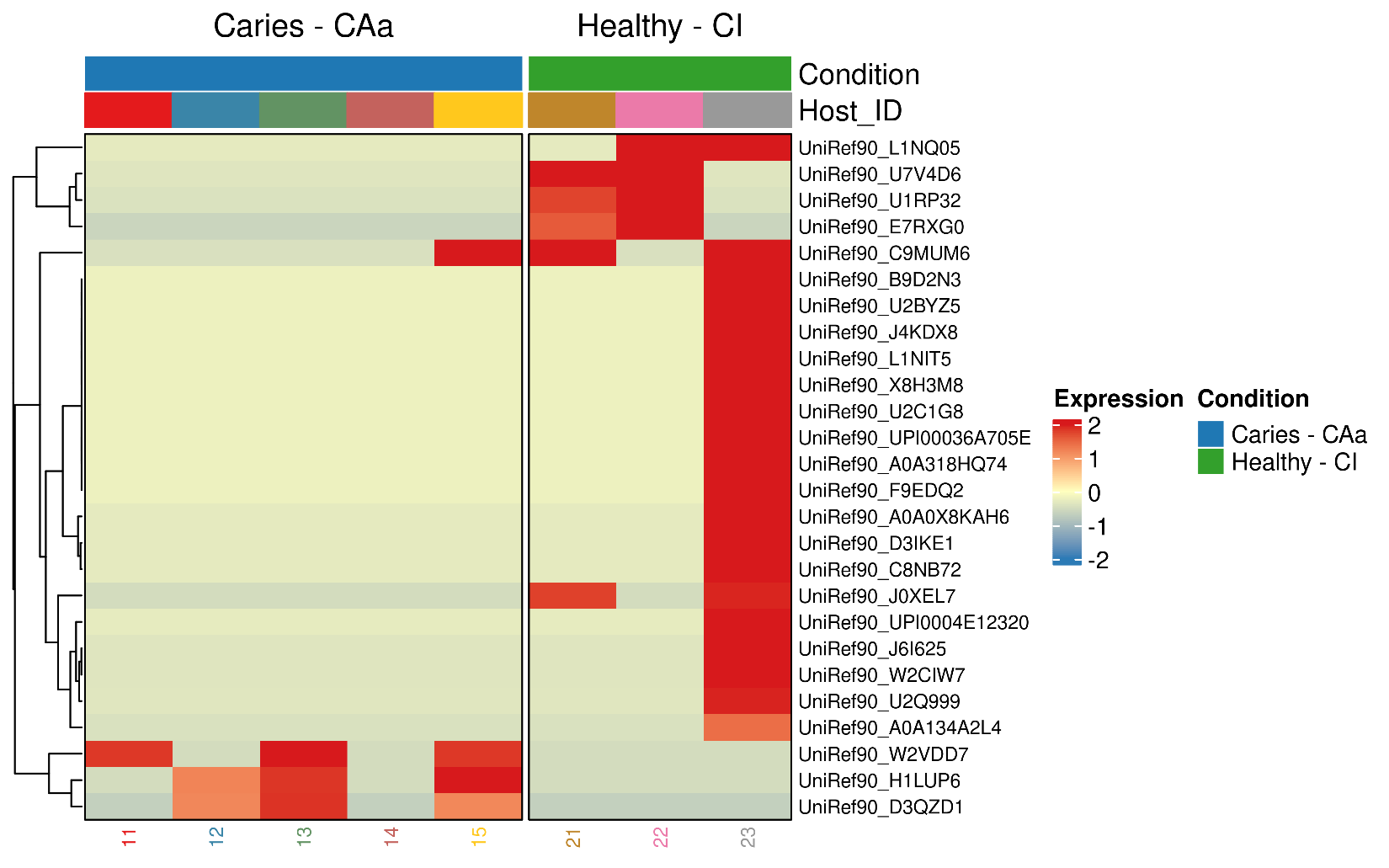

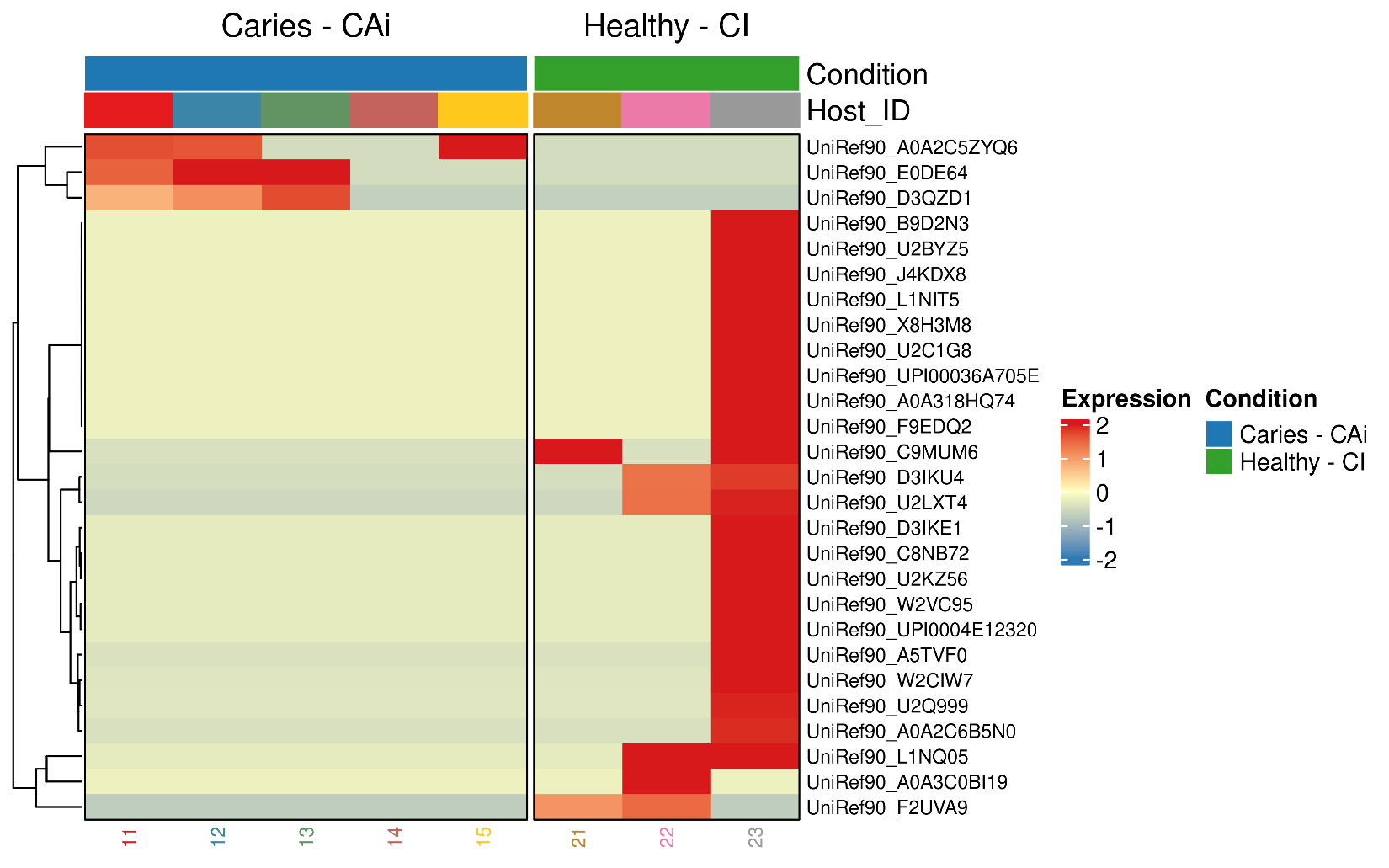

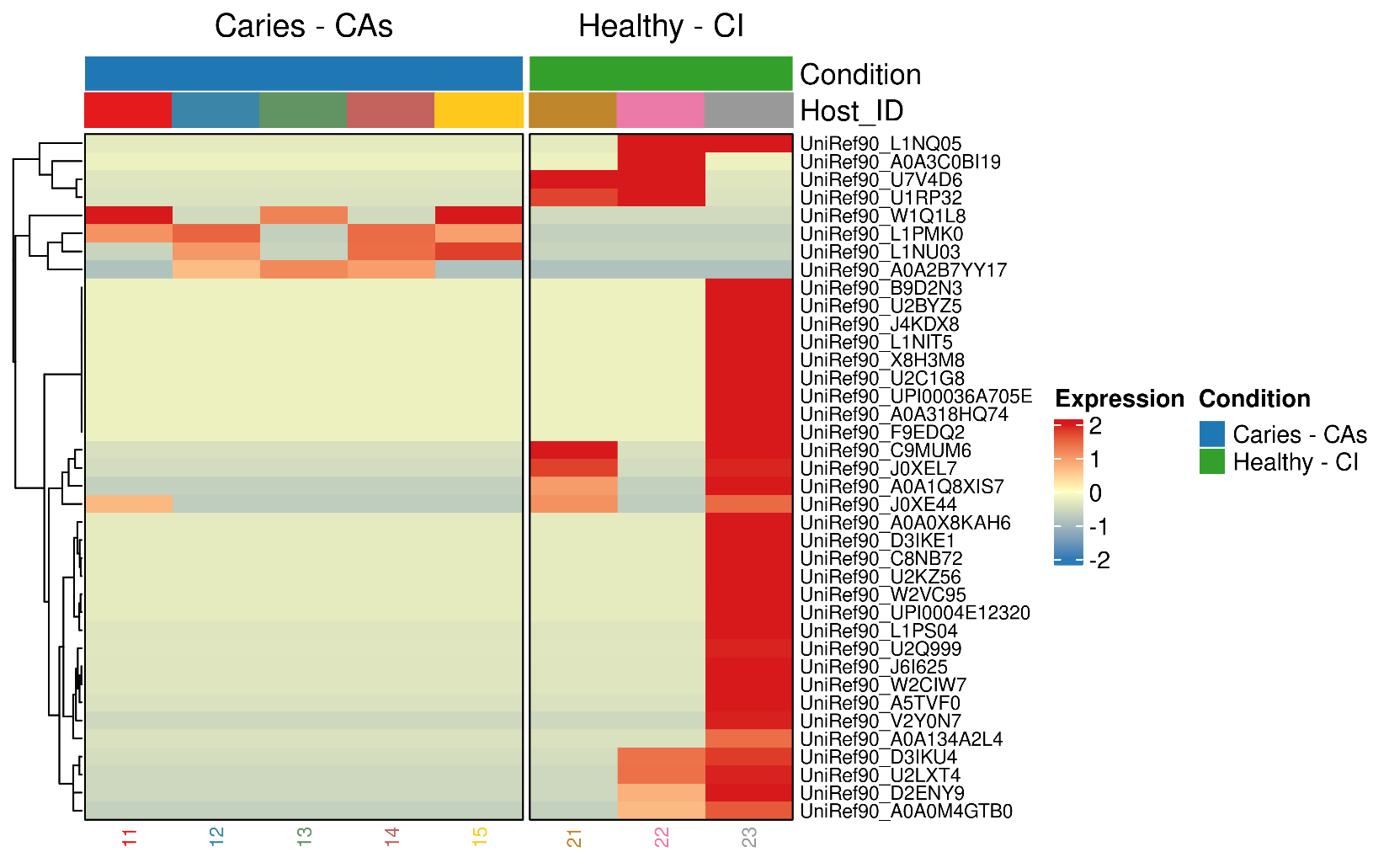

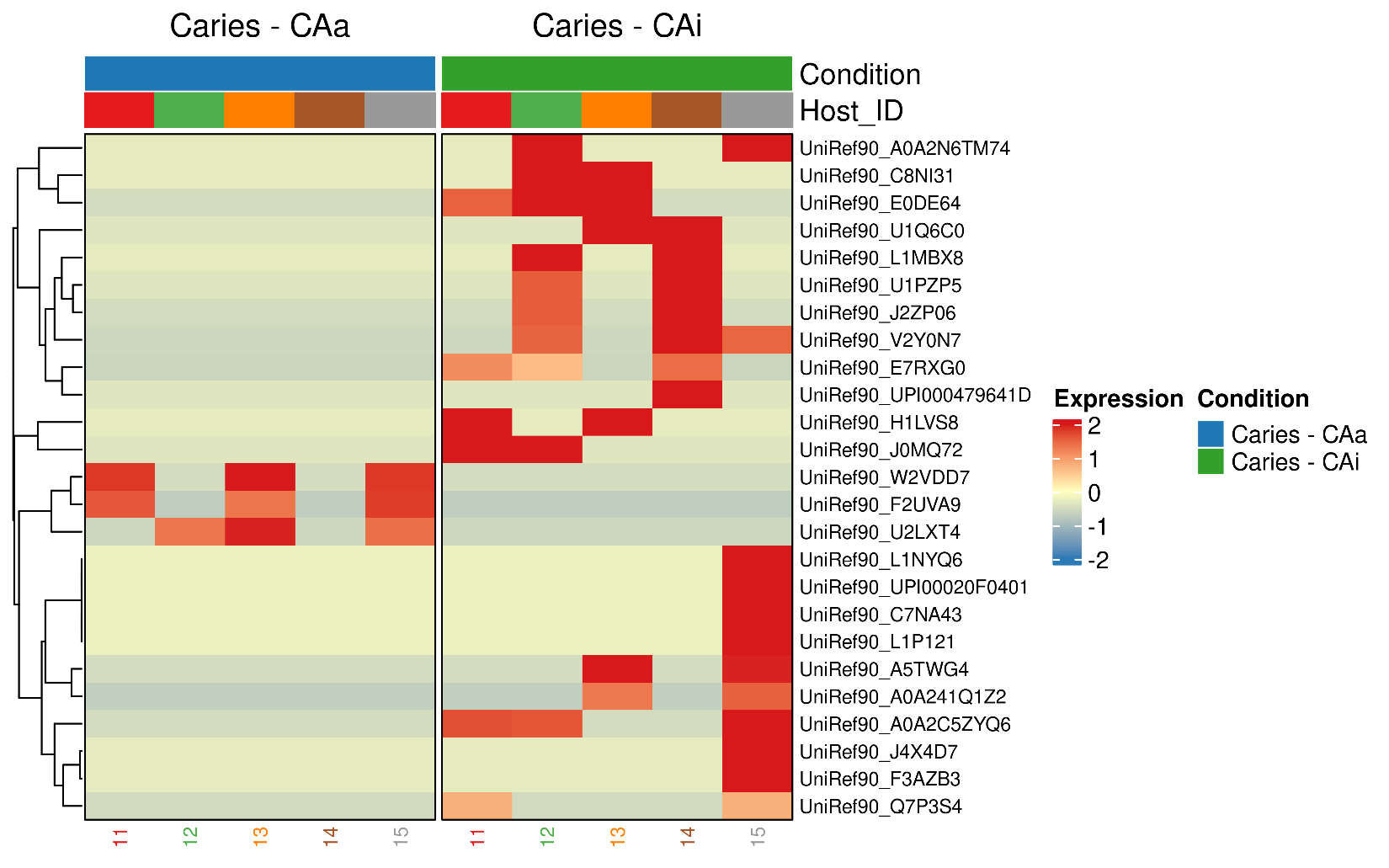

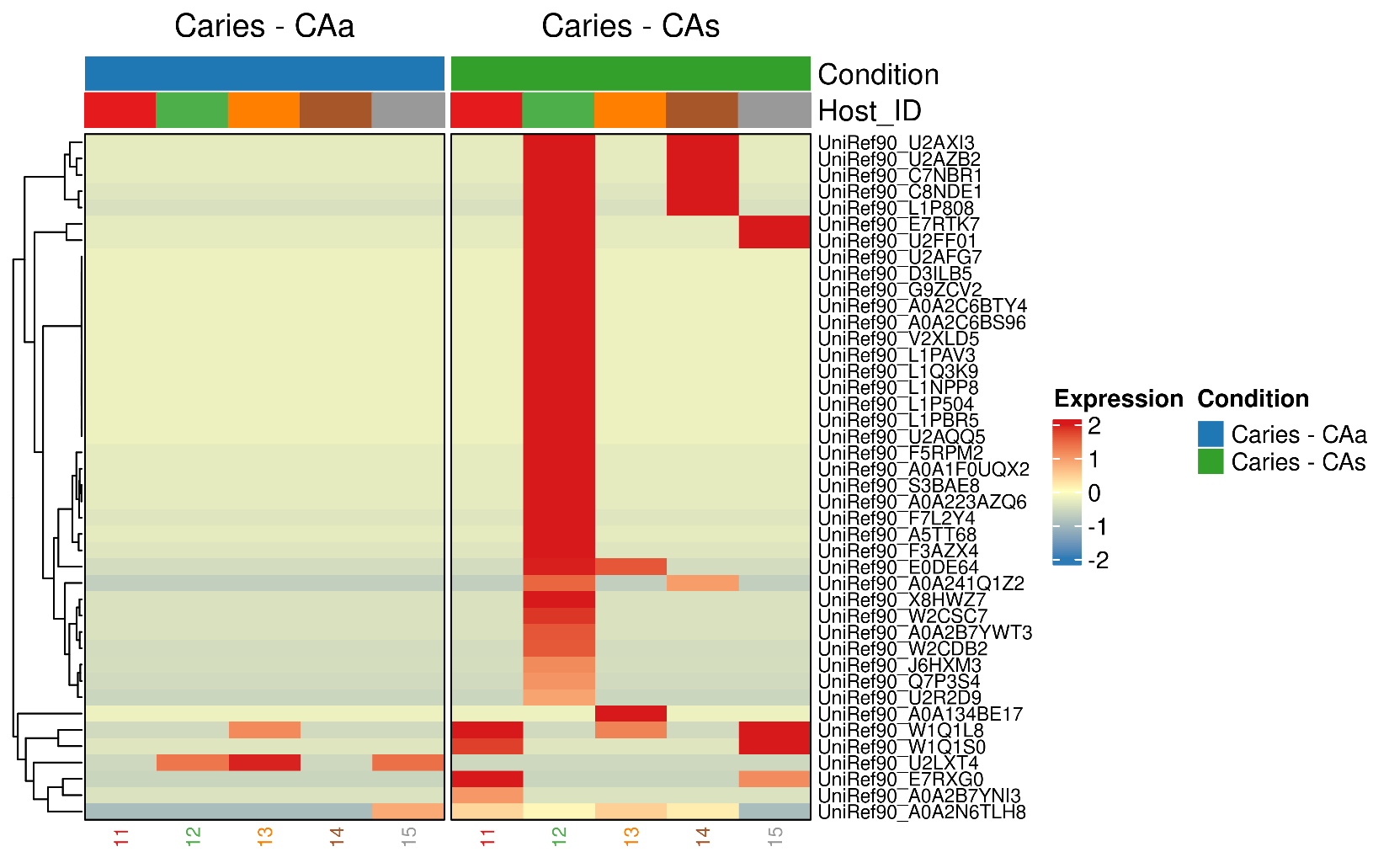

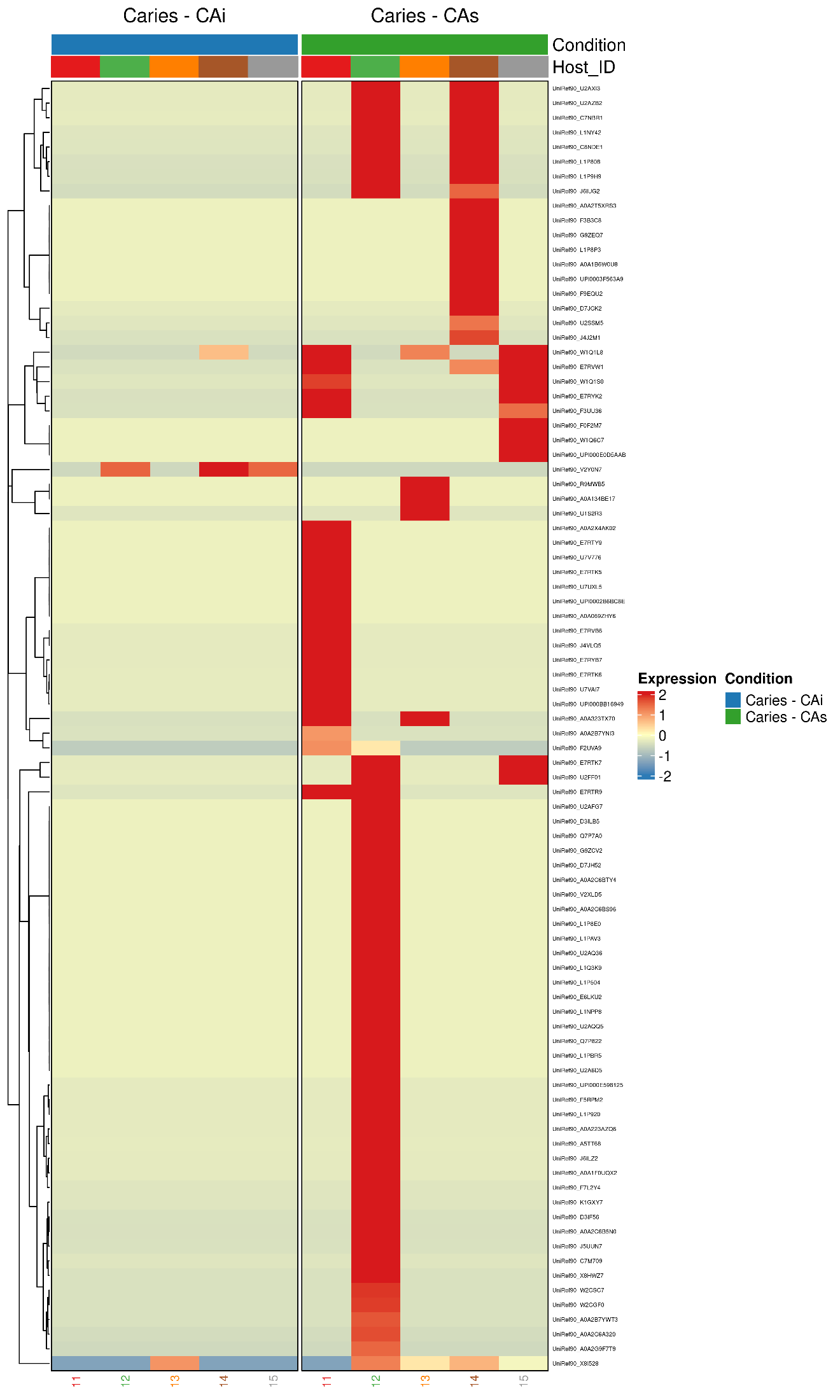
